# The genetic consequences of population marginality: a case study in maritime pine

**DOI:** 10.1101/2023.12.27.573436

**Authors:** Adélaïde Theraroz, Carlos Guadaño-Peyrot, Juliette Archambeau, Sara Pinosio, Francesca Bagnoli, Andrea Piotti, Camilla Avanzi, Giovanni G. Vendramin, Ricardo Alía, Delphine Grivet, Marjana Westergren, Santiago C. González-Martínez

## Abstract

**Aim:** Marginal tree populations, either those located at the edges of the species’ range or in suboptimal environments, are often a valuable genetic resource for biological conservation. However, there is a lack of knowledge about the genetic consequences of population’s marginality, estimated across entire species’ ranges. Our study addresses this gap by providing information about several genetic indicators and their variability in marginal and core populations identified using quantitative marginality indices.

**Location:** Southwestern Europe and North Africa.

**Methods:** Using 10,185 SNPs across 82 populations of maritime pine (*Pinus pinaster* Ait.), a widespread and economically important conifer characterised by a fragmented range, we modelled the relationship of seven genetic indicators potentially related to population evolutionary resilience, namely genetic diversity (based on all SNPs and two kind of outlier loci), inbreeding, genetic differentiation, recessive genetic load and genomic offset, with population geographical, demo-historical and ecological marginality (as estimated by nine quantitative indices). Models were constructed for both regional (introducing gene pool as random factor) and range-wide spatial scales.

**Results:** We showed a trend towards decreasing overall genetic diversity (albeit not necessarily for genetic diversity estimates based on outlier loci) and increasing differentiation with geographic marginality, supporting the centre-periphery hypothesis (CPH). Moreover, we found no correlation between population inbreeding and marginality, while geographically marginal populations had a lower recessive genetic load (only models without the gene pool effect) and ecologically marginal populations had a higher genomic offset, suggesting higher maladaptation to future climate.

**Main conclusions:** Overall, our results indicate that marginal populations could be at higher risk of maladaptation to climate change than core populations, despite reduced levels of genetic load, a risk that is exacerbated by typically small effective population sizes and increasing human impact.

## Introduction

All species thrive and reproduce within an environmentally limited geographic area, which sets the boundaries of their range. Current global warming may restrict the climatic suitability of some parts of the species’ ranges and, together with other factors (e.g., typically competitive biotic interactions, Loehle, 1998; but also positive, often unaccounted-for, interactions with other species, Stephan et al., 2021), modify their geographic and ecological margins. The ‘centre-periphery hypothesis’ (hereafter CPH), a major paradigm in biogeography that aims to disentangle the genetic, demographic, and ecological causes of species’ range limits (Gaston, 2009; Sexton et al., 2009), defines marginality as the level of geographic isolation from the species’ centre of distribution, which in turn is related to the species’ suitability for its environment (Brown, 1984; Hengeveld & Haeck, 1982). According to the CPH, marginal populations are expected to be less abundant and more prone to extinction than those in the centre, due to harsher environmental conditions at the periphery (Birch, 1957; Gaston, 2003; Nicholson, 1958; Richards, 1961; Whittaker, 1971). However, as specific environmental conditions in the core of the species distribution may also induce harsher environmental conditions than in the periphery (e.g., temperature extremes, rugged topography, peculiar edaphic features), Soulé (1973) distinguished between geographical and ecological marginality when describing centre-periphery gradients, defining ecological marginality as a population’s exposure to extreme environmental variables irrespectively of their geographical location.

While several studies have identified marginal populations based on different criteria (e.g., for maritime pine, Alía et al., 1996; Burban & Petit, 2003), we often lack evidence on the extent to which population marginality is associated with particular genetic features. Classic studies at the range-wide scale suggested a general trend for lower genetic diversity and higher genetic differentiation in marginal populations, but they typically did not distinguish between geographical and ecological marginality (Eckert et al., 2008; Johannesson & André, 2006; Pironon et al., 2015). More recent studies often called for decoupling the roles of geography and ecology in shaping range-wide patterns of genetic variation (Lira-Noriega & Manthey, 2014; Sexton et al., 2013). To that end, a clear distinction needs to be made between different kinds of population marginality. In a recent multispecies study, Picard et al. (2022) evaluated the ability of quantitative measures to distinguish between geographical, demo-historical (i.e., related to the demographic history of the target species) and ecological marginality.

Past global climate dynamics rather than demographic stochasticity seem to have played a crucial role in the establishment of current range limits (e.g., Hampe & Petit, 2005, for forest trees). Postglacial migrations after the Last Glacial Maximum (LGM) would have resulted in a mosaic of relatively small and isolated populations at the rear and, in a lesser extent, at the front edges of the expansion. In this context, the level of connectivity among marginal populations and between a given marginal population and the species distribution core is especially important (Sachdeva et al., 2022), as gene flow may increase genetic diversity and reduce differentiation (e.g., Lynch et al., 1995; Young et al., 1996). Gene flow can also bring adaptive alleles and contribute to the evolutionary rescue of small, isolated populations by buffering the effect of genetic drift, and reducing the fixation and accumulation of deleterious alleles within populations (Sachdeva et al., 2022). However, marginal populations may also carry specific alleles derived from local adaptation to atypical environments, constituting valuable genetic resources, and making the contribution of external gene flow harmful (i.e., the so-called ‘migration load’; Kimura et al., 1963).

Overall, marginal populations are considered to be more vulnerable to climate change than core populations (Kolzenburg, 2022; Soulé, 1973). Marginal populations are expected to accumulate deleterious variants (i.e., genetic load), a process governed by effective population size, *N*_e_ (Kimura & Ohta, 1969). However, genetic purging due to inbreeding tends to reduce genetic load over time, even in relatively small populations (Hedrick & Garcia-Dorado, 2016), and the overall outcome is context-dependant (Sachdeva et al., 2021). Current developments in population genomics have provided metrics to estimate maladaptation to future climates e.g., by estimating genomic ‘offsets’ or ‘gaps’ (a measure of the mismatch in genotype-climate association between current and potential future climates, Capblancq et al., 2020; Fitzpatrick & Keller, 2015; Gougherty et al., 2021; Rellstab et al., 2021). Studies aimed at validating genomic offset predictions with data from common garden experiments and natural populations have shown that populations with higher genomic offset exhibit a reduction in population growth performance (e.g., Fitzpatrick et al., 2021; for balsam poplar) and survival (Archambeau et al., 2024; for maritime pine), and concluded on the potential of this indicator to provide an estimate of the degree of expected maladaptation to future climate. Thus, despite several limitations (Rellstab et al., 2021; Láruson et al. 2022; Ahrens et al., 2023; Lotterhos, 2023), the calculation of genomic offsets may still enable for much needed systematic studies on the connection between population marginality and maladaptation in the face of climate change (Archambeau et al., 2024).

Maritime pine (*Pinus pinaster* Ait., Pinaceae) is an outcrossing, wind-pollinated conifer, with a widespread but fragmented natural distribution in southwestern Europe and North Africa, covering a wide range of contrasted environments from coastal dunes next to the Atlantic Ocean in France to the High Atlas Mountains in Morocco. Maritime pine population genetic structure is a consequence of historical and current dynamics of range expansion-contraction, resulting in distinct genetic clusters (Jaramillo-Correa et al., 2015). This species is also characterised by a fragmented distribution due to human-induced habitat loss and ecological disturbances such as forest fires (De-Lucas et al., 2009). Nowadays, some maritime pine populations are found in ecologically marginal environments (e.g., under very dry conditions in southern Spain and northern Morocco). In addition, the species’ range margins are characterised by small, geographically isolated populations, in particular in the southern and eastern parts of the distribution (Alía et al., 1996; Wahid et al., 2004). Reduced dispersal with distant core populations, coupled with demographic and environmental stochasticity, may push such populations into an ‘extinction vortex’ (Lande, 1988). Few studies have focused on describing the particular genetic characteristics of marginal populations of maritime pine (Salvador et al., 2000; Wahid et al., 2004; González-Martínez et al., 2007), whose potentially valuable genetic resources could be lost in the near future.

In this study, we used 10,185 Single Nucleotide Polymorphisms (SNPs) genotyped in 82 maritime pine populations (1,510 individuals), including ecologically and geographically marginal ones, to compute seven genetic indicators potentially related to population evolutionary resilience (defined as the property of an ecosystem to undergo adaptive evolution in response to biotic or abiotic disturbances; Sgrò et al., 2010) and correlated them with quantitative measures of marginality. The main objective of this study is to assess the relationship between marginality and population genetic features at the scale of the whole species range, by testing predictions of the CPH and adding new elements to its general framework. More specifically, we i) assessed the losses of genetic diversity and increases of genetic differentiation in marginal populations, distinguishing overall genetic diversity from that estimated using different kind of outlier loci; ii) evaluated the levels of accumulation of recessive genetic load based on counts of deleterious alleles; and iii) tested whether marginal populations are maladapted to future climate conditions, applying genomic approaches that consider the contribution of pre-adapted variants to future climates (i.e., genomic offset models).

## Materials and Methods

### Plant materials and molecular markers

Needles were collected from 1,510 individuals in 82 maritime pine populations covering all previously identified gene pools throughout the species range (Jaramillo-Correa et al., 2015; Milesi et al., 2023). Populations were selected based on gene pool size, density of maritime pine in the area, and representativeness in terms of climatic conditions specific to their geographical position (e.g., orography), as to achieve a regular sampling across the full distribution range of the species. This sampling is, to date, the most complete in the species (see Figure 1 and Table S1 in Supplementary Information), and includes several populations from the distribution margins, as well as isolated populations that have not been considered in genetic studies before. Population Cómpeta (COM), with only three samples, was removed from all data analyses but the gene-environment association (GEA) methods used to estimate the genomic offset. The accuracy of landscape genomic approaches, such as GEA, is highly improved by increasing the number of populations and environments, while being less sensitive to unbalanced sampling designs (Santos & Gaiotto, 2020). In addition, the two stands of Maures population (MAU) (see Table S1) were kept separate for these analyses as they were sampled at different altitudes.

**Figure 1.**
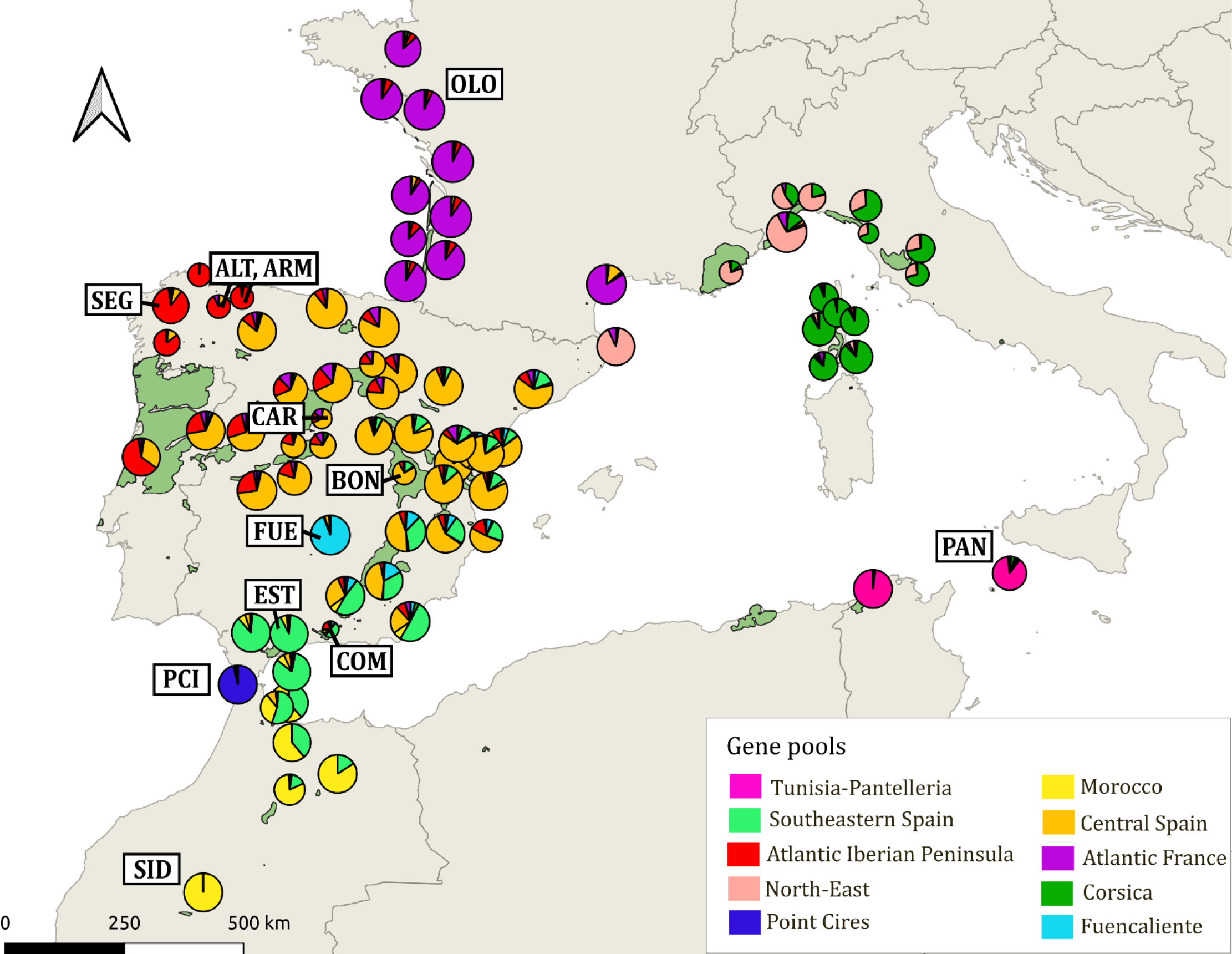
Genetic pools in *P. pinaster*. Pie charts depict membership proportions of each genetic cluster (*K*=10) for each studied population, calculated by STRUCTURE v2.3.4. The natural distribution of the species is shadowed in light green (see details in Figure S1). Population codes are only provided for those populations specifically mentioned in the main text (see full population information, including population names and geographical coordinates, in Table S1).

DNA was extracted using the DNeasy 96 plant kit (QIAGEN, Germany), following the manufacturer’s instructions. All the samples were genotyped for SNPs using the multispecies 4TREE Axiom array (Thermo Fisher Scientific, USA). For maritime pine, the 4TREE Axiom array combines SNPs identified in two previous studies: the 9K Illumina Infinium array generated by Plomion et al. (2016) and the exome capture experiment used in Milesi et al. (2023). The new array has a conversion rate of 79%, as well as 99% genotype reproducibility (based on genotyping of duplicated samples). Apart from potentially neutral genetic polymorphisms, this array comprises also SNPs from candidate genes that showed signatures of natural selection or significant environmental associations with climate at the range-wide spatial scale, orthologs for gene families with important adaptive functions in model species, and coding regions with differential expression under biotic and abiotic stress in maritime pine (see details in Plomion et al., 2016; Milesi et al., 2023). Only SNPs with high-quality scoring following the Best Practices Workflow implemented in the Axiom™ Analysis Suite v5.2 were selected and filtered by missing data (<30%), yielding a total of 10,185 SNPs. To assess the impact of missing data in the calculation of genetic indicators, SNPs were also filtered for missing data using a 5% threshold, resulting in a subset of 6,390 SNPs. SNP annotation based on SnpEff v5.1 (Cingolani et al., 2012) was retrieved from Cahn (2023) for a set of 1,325 SNPs in common with our study.

## Data analysis

### Population marginality indices

To assess population marginality, we first produced a new distribution map for maritime pine that includes only natural populations (Figure S1), building on that of Caudullo et al. (2017) but adding information from National Forest Inventories and specific publications for less-known parts of the distribution range (Alía et al., 1996; Marques et al., 2012; Abad-Viñas et al., 2016; Fkiri et al., 2019; Wahid et al., 2004, 2006). Second, we computed eight quantitative marginality indices that consider both geographical distribution and demographic history, following Picard et al. (2022) (see Table 1 and Table S2 in Supplementary Information). The demo-historical indices, which are related to the postglacial colonization history of the species (see, e.g., Bucci et al., 2007; Jaramillo-Correa et al., 2015), and the centroid index were computed from the distribution map. The centroid of the species’ distribution corresponded to the location whose geographic coordinates were the average of the geographic coordinates of all locations where the species is present (in our case, eastern Spain, a known glacial refugia of maritime pine; Salvador et al., 2000; see Figure S2). Then, the centroid index was defined as the ‘cost distance’ between any population and the centroid location. Cost distances were computed using a conductance matrix (the inverse of a resistance matrix), reflecting the conductance of gene flow. In the conductance matrix, sea cells were assigned low conductance, land cells where the species was absent were assigned intermediate conductance, and land cells where the species was present were assigned high conductance (see Table 1). This index reflects the level of long-distance gene flow between a given population and the geographical core of the species distribution. The other geographical indices relied on a morphological spatial pattern analysis (MSPA), which considers a binary image (1-presence/0-absence and NA for water) with emphasis on the connectivity within the image (Soille & Vogt, 2009). Presence/absence maps were structured in three categories (cores, edges, and other classes including loops, islets, bridges and branches) using BioManager package in R version 4.2.2. Third, we calculated an index of ecological marginality (henceforth ‘ecological index’) based on climatic data for the period 1901-1970, as follows. For each population, we extracted the climatic information provided by the Climate Downscaling Tool (ClimateDT, https://www.ibbr.cnr.it/climate-dt/) for the Summer Heat Moisture (SHM) aridity index, an indicator of exposure to drought (see, e.g., De La Torre et al., 2014; Marchi et al., 2020), and temperature (bio4) and precipitation seasonality (bio15). This set of climatic variables was selected because it best explained the climatic variation across the species range in previous studies (e.g., Archambeau et al., 2024). The ecological index was constructed by computing the standardized Euclidean distance for SHM, bio4 and bio15 between each population and the overall average. Thus, this index represents the climatic distance of the population from the average climate (Table S2). Finally, we reduced the set of marginality indices by removing those that were highly correlated based on Pearson’s correlation coefficients (Pearson’s correlation ≥ 0.6) and a principal component analysis (PCA) using *FactoMineR* package in R v4.2.2 (Figure S3).

**Table 1.** Geographical (*geo*) and demo-historical (*histo*) marginality indices computed for maritime pine populations; ‘(-)’ and ‘(+)’ indicate that higher values of the index correspond to lower and higher population marginality, respectively; ‘(-)(+)’ indicates that both higher or lower values of the index correspond to higher population marginality. Adapted after Picard et al. (2022).

| Marginality index | Category | Link with marginality | Description |
| --- | --- | --- | --- |
| Area | *geo* | (-) | Size in ha of the core (i.e., continuous patches of the species’ presence) that is the nearest to the location of the studied population. |
| Gravity | *geo* | (-) | Spatially continuous alternative to the area index, computed as the weighted mean of the core areas with weights inversely proportional to the distance squared from the location to the cores. |
| Centroid | *geo* | (+) | Eccentricity computed as the cost distance from the centroid of the species distribution to any location, taking into account the terrestrial and marine connectivity linked to pollen circulation capacity. Cost distances are based on a conductance matrix (the inverse of a resistance matrix) reflecting the permeability of land and water cells to gene flow. This index is expected to capture the level of long-distance gene flow. |
| Edge | *geo* | (-) | Distance to the nearest border of the species distribution. |
| Isolation | *geo* | (+) | Isolation with respect to the core species distribution, computed as the distance from the focal population to the nearest core greater than 100 ha and further than 50 km away. |
| Second nearest-core | *geo* | (+) | Spatially continuous alternative to the isolation index, computed as the distance from a location to the second nearest-core greater than 100 ha. |
| North-South | *histo* | (-) (+) | Proximity to the species rear-edge (southernmost limit of the species distribution) or leading-edge (northernmost limit of the species distribution) along a latitudinal gradient. |
| East-West | *histo* | (-) (+) | Proximity to the species leading-edge or rear-edge along a longitudinal gradient. |

### Population genetic structure and genetic indicators

Population genetic structure was evaluated using the Bayesian clustering algorithm implemented in STRUCTURE v2.3.4 (Pritchard et al., 2000). To assess the optimal individual’s assignment probabilities (*Q_ancestry_*) in *K* genetic clusters (or gene pools), we ran models from *K*=1 (no structure) to *K*=10 with a burn-in length of 100,000 and run lengths of 200,000 MCMC steps. The number of *K* that best describes the genetic structure was determined based on the delta *K* method (Evanno et al., 2005) and the visual observation of bar plots.

All genetic indicators were computed at the population level using the full dataset of 10,185 SNPs (Table S3). In a few cases where more than one stand was sampled for a population (see Table S1), genetic indicator values were averaged. The computation of genetic indicators was robust to the inclusion of missing data, as shown by the high correlations (Pearson’s correlation > 0.7) with genetic indicator estimates computed using the dataset (6,390 SNPs) with missing data lower than 5% (Figure S4a,b).

Genetic diversity was estimated as 1-*Q_inter_*, with *Q_inter_* being the observed frequencies of identical pairs of alleles among individuals within populations, using GenePop v4.7.5 (Rousset, 2008), after standard correction for sample size (N), using (N/N-1). Notice that this estimate averages over monomorphic and polymorphic loci. In addition to overall genetic diversity, we also calculated two indicators of genetic diversity based on outlier loci. The first was calculated on the basis of climate-associated (GEA) outliers (i.e., 73 outlier SNPs selected for the genomic offset computation, described below). The second indicator was calculated on the basis of general outliers due to unknown factors (i.e., 151 outlier SNPs common to two environment-independent *F*_ST_-outlier-detection methods, as implemented in the R package *pcadapt* and BayeScan v2.1; Foll & Gaggiotti, 2008). Population inbreeding (*F*_IS_) was estimated following Weir & Cockerham (1984). Population-specific divergence was estimated as the genetic differentiation (population-specific *F*_ST_) of each population from a common ancestral gene pool using BayeScan v2.1 (Foll & Gaggiotti, 2008), as well as Jost’s *D* (Jost, 2008) estimated with MMOD package in R v4.2.2.

Finally, we computed two additional genetic indicators more specifically related to potential maladaptation to climate change, i.e., the recessive genetic load and the genomic offset. The recessive genetic load represents the accumulation of predicted deleterious mutations in the population standardized by the population genetic diversity. This statistic was estimated by counting different kind of mutations (annotated by SnpEff v5.1; Cingolani et al., 2012) averaged over individuals, as the number of derived moderate-(i.e., non-synonymous) and high-impact (i.e., loss of function) mutations in homozygosity divided by the number of derived low-impact (i.e., synonymous) mutations in homozygosity, following González-Martínez et al. (2017). The genomic offset is estimated as the change in genetic composition required to maintain the current gene-climate relationships under future climates (see Fitzpatrick & Keller, 2015) and thus captures the degree of maladaptation a population will undergo when the environment to which it is currently adapted will change, either from a spatial or temporal perspective (Rellstab et al., 2021; Lotterhos, 2024). Briefly, we first identified outlier SNPs for climate adaptation with two univariate GEA methods, BAYPASS (Gautier, 2015) and Latent Factor Mixed Model (LFMM; Frichot et al., 2013), and three multivariate ones, Redundancy Analysis (RDA; Capblancq et al., 2020), partial RDA (Van Den Wollenberg, 1977) and Gradient Forest (GF; Ellis et al., 2012). Then, the genomic offset was estimated using the set of outlier SNPs identified by at least two methods (see the GitHub detailing this analysis and referenced below) and the GF approach, which showed the best empirical validation in a previous study based on a smaller sample of populations and SNPs (Archambeau et al., 2024), and six climatic variables related to maritime pine expected exposure to climate change (Table S4; see also Archambeau et al., 2024). Future climates for 2070 were described using the predictions from the moderately alarming shared socio-economic pathway (SSP) SSP3-7.0 and five global circulation models (GCMs; IPCC, 2021). As genomic offset predictions across GCMs were highly correlated (Pearson’s correlation coefficient >0.75; Figure S5), we used population averages for the five GCMs. Additional details and scripts are available at https://anonymous.4open.science/r/ReadyToGO_Pinpin-FA56/README.md. All analyses were undertaken under R v4.2.2 (R Core Team, 2022).

### Effects of population marginality on genetic indicators

We estimated the relationship between genetic indicators and population marginality by fitting two series of seven linear mixed-models with pairwise interaction terms (one model for each genetic indicator), using the R packages LME4 and LMER, respectively. Models M1 to M7 included population marginality indices as fixed effects irrespectively of the gene pool of origin while models M8 to M14 also included the gene pool of origin as random effects. Random effects in linear mixed-models allow the inclusion of non-independent data from a nested structure (populations sampled within gene pools), allowing each level of the grouping factor (gene pool) to have its own random intercept. The gene pools with a single population (Fuencaliente, FUE; and Point Cires, PCI) were assigned to the geographically closest gene pool (Southeastern Spain and Morocco, respectively). Model goodness-of-fit was evaluated with *R*² and both the Akaike Information Criterion (AIC) and the Bayesian Information Criterion (BIC). A visual evaluation of model fit to data was also performed using diagnostic plots (QQ and residual plots; Figures S12 to S17). Then, the best models were selected by considering goodness-of-fit (higher *R*² and lower AIC/BIC) and parsimony criteria (i.e., including only significant effects at α = 0.01).

## Results

### Marginality indices

Pairwise Pearson’s correlations and PCA identified some strongly correlated marginality indices (Figure S3). For example, the second nearest-core was positively correlated with the isolation index (Pearson’s correlation of 0.64) and negatively correlated with the edge index (Pearson’s correlation of -0.67), and the ecological index was negatively correlated with the East-West index (Pearson’s correlation of -0.51). Thus, only five indices with low correlation (<0.6) were retained for further analysis, namely three geographical indices (centroid, second nearest-core and gravity; see definitions in Table 1), one demo-historical (North-South), and the ecological index based on climate distances (Figure S6). Interestingly, the centroid index was positively correlated with the longitude (Pearson’s correlation of 0.70) and the North-South index was negatively correlated with the elevation (Pearson’s correlation of -0.56), with southern populations being, generally, at higher elevation (Figure S3).

### Population genetic structure

Population genetic structure analyses identified ten distinct gene pools, among which two included only a single population (FUE in southern Spain and PCI in northern Morocco; Figure 1). Remarkably, these two single-population gene pools were not identified as marginal populations by the geographical or demo-historical marginality indices whereas one of them, PCI, was characterized by high values of the ecological index (standardized value of 5.013; Table S2), indicating persistence in a marginal climate. FUE and PCI had also low levels of admixture with nearby gene pools. In contrast, the eight main gene pools (with the exception of the highly isolated Tunisia-Pantelleria one, see below) were not genetically isolated from each other, with populations often showing admixture with nearby gene pools. This suggests either historical or recent gene flow across neighbouring gene pools along a latitudinal cline in the western range of the species, and substantial shared ancestry among French (including Corsica) and Italian populations in the eastern one.

### Effects of population marginality on genetic indicators

Our models revealed a decline in overall genetic diversity (corrected 1-*Q_inter_*, Figure 2a) and an increase in genetic differentiation (population-specific *F*_ST_, Figure 2b; and Jost’s *D* statistic, Table S5) with population marginality based on the centroid, second-nearest core and North-South indices (Tables 2a and S4). Indeed, models M1 and M5 predicted lower genetic diversity and higher genetic differentiation in marginal populations based on the centroid index (Tables 2a and S4), in particular in isolated populations from the southern maritime pine range (as shown by significant coefficients for the second nearest-core and North-South indices and their interaction; see also Figures 2 and S7). Model M2 revealed a decrease in genetic diversity based on GEA outliers for southern range populations (Table 2a; Figure S8a). Interestingly, model M3 revealed an increase in genetic diversity based on general outliers in marginal northern range populations, especially for those with high values of the centroid index (as shown by significant coefficients for North-South and centroid indices and their interaction; Table 2a, Figure S9). However, this unexpected pattern may just result from confounded effects due to high genetic diversity for general outlier loci in the two northernmost gene pools (Atlantic France and North-East; compare Figure 1 with Figure S8b and see M10 below). Models including the gene pool of origin as a random effect (M8 and M12; Tables 2b) significantly improved the fitting by 10% for overall genetic diversity and 20% for population-specific *F*_ST_ (29% for Jost’s *D* statistic; Table S5), but retained only the centroid and North-South indices and their interaction as explanatory factors (with the interaction having a similar interpretation as that for models without gene pool as random effect; Figure S10). For genetic diversity based on GEA outliers, the model including the gene pool as a random effect (M9) also improved the fitting (by over 30%) and revealed the same relationship with the North-South index as M2 while no significant relationship was found between genetic diversity based on general outliers and population marginality at the gene pool level (M10). Interestingly, we found no relationship between inbreeding and the indices of population marginality (see Figure S8c) for any model (M4, M11; Table 2a,b).

**Figure 2.**
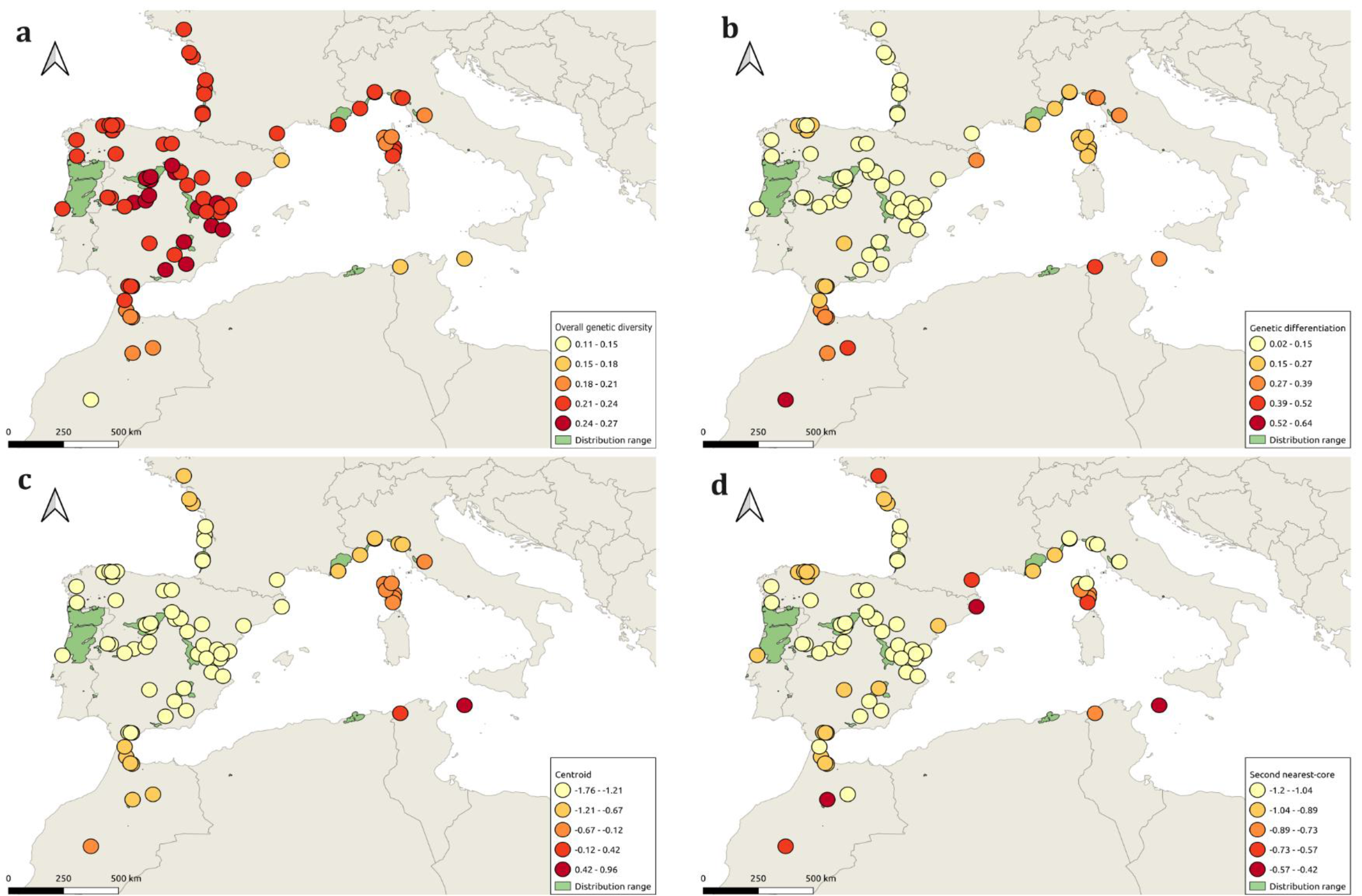
Geographical distribution of a) overall genetic diversity (1-*Q_inter_*) and b) genetic differentiation (population-specific *F*_ST_), and two marginality indices involved in significant correlations with these genetic indicators, c) centroid index and d) second nearest-core index, for *P. pinaster* populations. See Table S1 for population information, including population names and geographical coordinates.

**Table 2.**
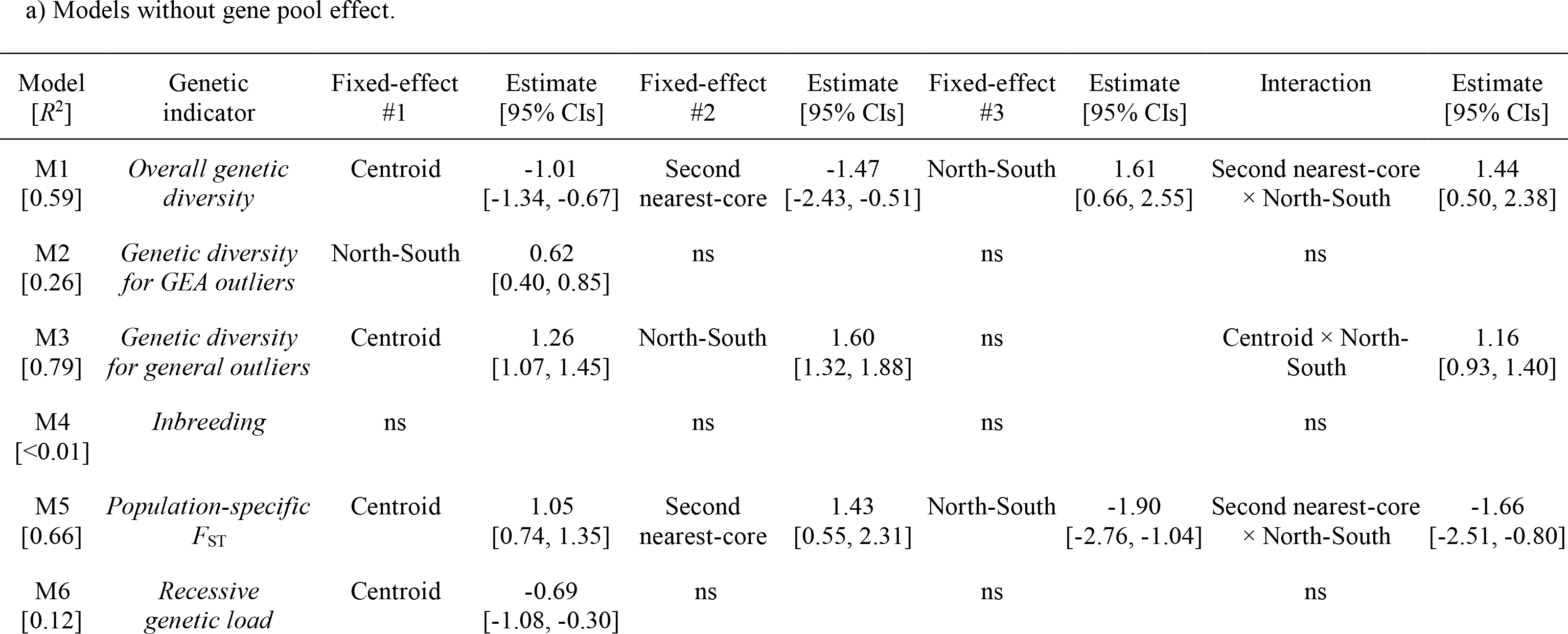
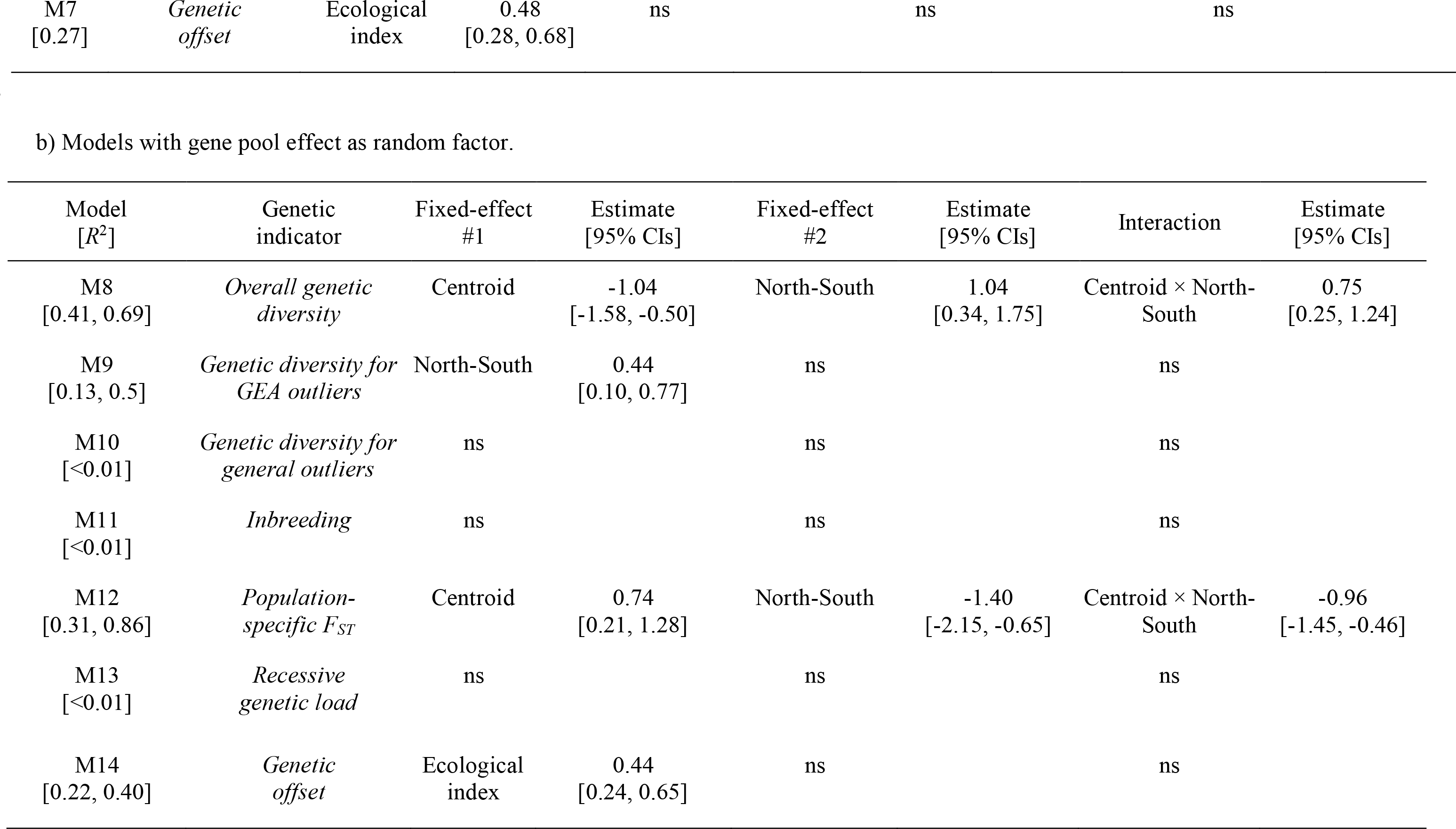
Models evaluating the effect of population marginality on the estimated genetic indicators; a) Models without gene pool eff ect (M1-M7); b) Models including gene pool effect as a random factor (M8-M14). Significant fixed effect(s) are listed with their associated point estimates and 95% confidence intervals [in brackets]. *R*^2^: variance explained by fixed factors (M1-M7) or both fixed (marginal *R*^2^) and random (conditional *R*^2^) factors (M8-M14); ns: not significant.

The model for recessive genetic load (M6) showed reduced genetic load for marginal populations based on the centroid index, this association probably stemming mainly from the high recessive genetic load found in some Iberian core populations (e.g., Carbonero el Mayor, CAR, and Boniches, BON; see Figure S8d). However, this model had a relatively low goodness-of-fit (*R*² = 12%) and we found no relationship between recessive genetic load and any of the population marginality indices when we added the gene pool of origin as a random effect (M13, Table 2b). Temperature seasonality (bio4) was the most important predictor contributing to the genetic turnover in genomic offset estimates (see Figure S11b and the GitHub detailing this analysis). Noticeably, this variable showed a steep slope between -2 and 0°C, which may indicate a rapid turnover in allele frequency in this range (see Figure S11a). Despite relatively poor goodness-of-fit (Table 2a,b and Figures S16 and S17), both models including the genomic offset, without (M7) and with (M14) the gene pool of origin as a random effect, predicted an increased genomic offset for populations in marginal climatic conditions (Table 2a,b and Figure 3b). Accordingly, we observed a trend for higher genomic offset in the western gene pools along the Atlantic coast (average genomic offset of 0.047 ± 0.026 and 0.047 ± 0.016 for the Atlantic Iberian Peninsula and the French Atlantic gene pools, respectively), with remarkably high values for Segurde (SEG), Alto de la Llama (ALT), Armayán (ARM) and Olonne sur Mer (OLO) as well as for some southern Mediterranean populations (e.g., Point Cires, PCI; Fuencaliente, FUE; Estepona, EST; and Pantelleria, PAN; Figure 3a).

**Figure 3.**
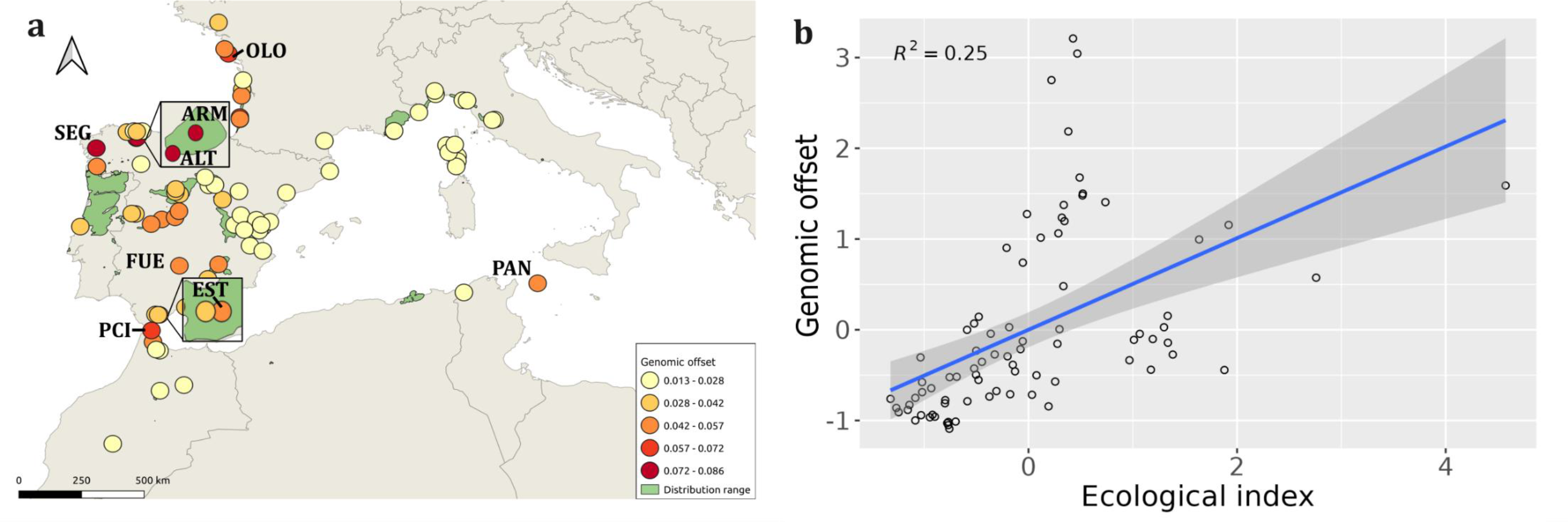
a) Geographical distribution of genomic offset for *P. pinaster* populations and b) correlation between the genomic offset and the ecological index based on climate distances (standardized values), as shown by linear regression in *ggplot2* R package. Population codes are only provided for populations specifically mentioned in the main text (see full population information, including population names and geographical coordinates, in Table S1).

## Discussion

In this study, we revealed a trend towards declining genetic diversity and increased genetic differentiation in geographically and demo-historically marginal populations of maritime pine; however, this genetic diversity trend would not necessarily apply to genes potentially involved in local adaptation, as shown by genetic diversity analyses based on different sets of outlier loci. We also found lower recessive genetic load in geographically marginal populations and higher genomic offset in ecologically marginal ones. Models including the gene pool of origin as a random effect were similar to those without (with the notable exception of the models for genetic diversity for general outliers and recessive genetic load), suggesting that the underlying processes operate at both the regional and range wide geographical scales in this species. These results, taken together, provide support for the CPH and suggest that climate change may endanger valuable and untapped genetic resources in maritime pine.

### Patterns of genetic diversity and differentiation support the CPH

The lower overall genetic diversity and higher genetic differentiation of geographical and demo-historical marginal populations support the main predictions of the CPH. Our results are consistent with those of Eckert et al. (2008) and Pironon et al. (2016), who found a decline in genetic diversity and an increase in differentiation towards the limits of the species ranges in 47% and 45% of studies in various taxa, respectively. As in our study, lower genetic variation in peripheral compared to central populations was found for some other conifers, i.e., in Sitka spruce (*Picea sitchensis*; Gapare et al., 2005), Swiss stone pine (*Pinus cembra*; Gugerli et al., 2009) and common yew (*Taxus baccata*; Hilfiker et al., 2004), but not in Norway spruce (*Picea abies*), for which only genetic differentiation conformed to the CPH (Westergren et al., 2018). Interestingly, unlike the distribution of overall and climate-associated (GEA) outlier genetic diversity with population marginality, genetic diversity based on general outliers (identified by the R package *pcadapt* and BayeScan v2.1; see Methods) increased with geographic marginality in terms of distance from the centroid, especially for populations in the northern margins. There are several ways to interpret this result. Firstly, geographically marginal populations may show more variation in outlier SNPs linked to unknown biotic or abiotic factors (other than climate) than central populations. Secondly, this pattern could result from a confounding effect linked to the distinct level of genetic variation in outlier SNPs in the different gene pools, which could be corroborated by the fact that no significant relationship with population marginality remains when models included the gene pool of origin as a random factor. Finally, we could hypothesize that no robust biological explanation can be drawn from this result, due to the uncertainty of these outliers to be related or not to adaptation. Indeed, Lotterhos et al. (2014, 2015) and Hoban et al. (2016) showed that most outliers detected with *F*_ST_-outlier tests are likely to be false positives when calculated on species with peculiar demographic histories such as range expansion (e.g., Travis et al. 2007; Excoffier et al. 2009), as is the case in maritime pine.

In addition to geographical marginality, demo-historical marginality showed a significant association with overall genetic diversity (M1 in Table 2a), as well as with genetic diversity based on outlier loci (M2 and M3). This significant association is consistent with the pattern observed in plants in the Mediterranean basin (Fady & Conord, 2010) but differs from a more global pattern in which the neutral genetic diversity of plants (including pines) does not decrease significantly with latitude (De Kort et al., 2021). Genetic differentiation, which decreases with latitude in plants across their ranges (Gamba & Muchhala, 2020), showed a significant association with demo-historical marginality in the same direction as well (see M5 in Table 2a and M5bis in Table S5). However, despite the general trend of decreasing genetic diversity with latitude, southern populations from the Moroccan and Tunisia-Pantelleria gene pools and northern populations from the North-East and Corsican gene pools exhibited the lowest overall and GEA outlier genetic diversity, and the highest genetic differentiation in maritime pine (see Figures 2 and S8a). Maritime pine gene pools are probably the result of population expansion from multiple glacial refugia, both in Mediterranean and Atlantic regions of the species (Bucci et al., 2007; Jaramillo-Correa et al., 2015), with Naydenov et al. (2014) also suggesting that, for this species, the high level of overall genetic differentiation may have resulted from long historical isolation predating the Last Glacial Maximum (∼18,000 years ago). This was confirmed by estimates of genetic divergence between North-African (Moroccan) and Iberian populations of maritime pine, which dated back to 1.90 Ma (95% credibility interval: 1.41-2.76), probably due to the Strait of Gibraltar’s effect as a major biogeographic barrier to pollen and seed gene flow (Jaramillo-Correa et al., 2010). Consequently, southernmost Moroccan populations are likely to have been pre-glacial relict populations that survived in North-African refugia (Baradat & Marpeau, 1988; Vendramin et al., 1998). After range expansion from glacial refugia, the persistence of maritime pine in several isolated groups characterized by contrasting climatic conditions may have resulted in populations that are locally adapted to the climate in some way. However, our models revealed that the southernmost populations of this species (Moroccan and Tunisia-Pantelleria gene pools) may be more at risk of not displaying enough diversity, neutral or adaptive, than the central populations. This pattern may have been exacerbated by a reduction of effective population size (*N*

) due to human impact (Wahid et al., 2004), as well as the incidence of recurrent forest fires resulting in population bottlenecks and genetic drift in this region (Vendramin et al., 1998). The demographic history of maritime pine in the north-eastern part of the continental range and the Corsica island has not been clearly assessed (Naydenov et al., 2014), however, the presence of an endemic mitotype in this region suggests long-term isolation of North-East and Corsican gene pools (Burban & Petit, 2003). Nevertheless, both overall and GEA outlier genetic diversity for these populations seem to be larger than for those from the southernmost gene pools (see Figures 2 and S8a).

### Lower recessive genetic load in geographically marginal populations

Our models revealed a significant, albeit weak, reduction in recessive genetic load with increased geographical population marginality. However, these models were only significant when the gene pool effect was not accounted for, suggesting the existence of gene pools with reduced/increased recessive genetic load and an important role of demographic history (see below). This was a surprising finding, as we were expecting marginal populations to be characterised by an accumulation of recessive genetic load due to the reduced effectiveness of purifying selection in small and isolated populations with high demographic stochasticity (Caballero et al., 2017; Sachdeva et al., 2022). The level of accumulation of deleterious mutations and the extent to which it represents a risk to a given population depends primarily on its effective size (*N*_E)_. Previous empirical studies of animal and plant populations that underwent historical range expansions or declines have often shown an increase in genetic load (e.g., Günther & Schmid, 2010, in *Arabidopsis thaliana*; González-Martínez et al., 2017, in *Mercurialis annua*; Feng et al., 2019, in *Nipponia nippon*; or Peischl et al., 2013, in humans). However, as in maritime pine, recent empirical studies based on genomic data suggested that recessive genetic load can also be purged in long-term isolated and inbred populations (see review in Dussex et al., 2023). As an example, Dussex et al. (2021) found that current island populations of *Strigops habroptilus*, a New-Zealand flightless parrot, had lower deleterious mutation load compared to mainland populations. A similar pattern was found for the Alpine ibex (*Capra ibex*), which suffered severe population bottlenecks and nearly became extinct (Grossen et al., 2020), and for the Channel Island fox (*Urocyon littoralis*) that has remained at small population sizes with low diversity for many generations (Robinson et al., 2018).

Present-day levels of inbreeding in maritime pine are low and not significantly higher in marginal than core populations. Therefore, we hypothesise maritime pine marginal populations to have effectively purged recessive genetic load during past inbreeding events, operating at the regional scale. These events may have occurred during the range contractions and/or expansions associated to Quaternary glacial and interglacial forest tree migrations (Bucci et al., 2007; Naydenov et al. 2014; Jaramillo-Correa et al., 2015). Polyembryony, which is ubiquitous in gymnosperms such as maritime pine (Willson & Burley, 1983), could have also played a role in purging recessive genetic load, as it tends to dampen self-fertilisation’s deleterious effects by more effectively removing the mutational load through selection between viable embryos (see Latta, 1995).

### Higher genomic offset in ecologically marginal populations

Our models also revealed a higher genomic offset for ecologically marginal populations, but not for geographically or demo-historically marginal ones suggesting that the gap between the current and required genetic composition in future climates will be greater for populations in marginal climatic conditions. This is consistent with the findings of Fréjaville et al. (2019), who observed that adaptation lags in several forest trees, including maritime pine, are consistently higher in climatically (cold/warm, dry/wet) marginal populations than in populations growing under climatically optimal conditions. Although studies estimating genomic offset in marginal populations are still scarce, two recent studies in widespread Asian forest trees provided support to our findings, as they showed a relatively high genomic offset in the northern and southern distribution margins of the Chinese thuja tree (*Platycladus orientalis*; Jia et al., 2020) and the sawtooth oak (*Quercus acutissima*; Yuan et al., 2023); but whether these range margins represented also ecologically extreme environments was not assessed and more detailed studies are thus needed.

Estimation of the genomic offset is becoming a popular approach to assess population vulnerability in the face of climate change. Genomic offset predictions have been validated using data from common garden experiments and natural populations (e.g., Fitzpatrick et al., 2021), including for maritime pine (Archambeau et al., 2024), but it is not free of pitfalls (see Ahrens et al., 2023; Rellstab et al., 2021; Lind et al., 2024; Archambeau et al., 2024).

Moreover, genomic offsets can gauge (at some extent) for maladaptation to future climates but not for the adaptive capacity of populations (Fitzpatrick et al., 2021; Archambeau et al., 2024). Despite some of our model predictions based on general outlier loci, which showed a reduction in potentially adaptive diversity towards the distribution margins, marginal populations of forest trees can retain notable adaptive capacity, as shown in common garden experiments for the handful of species with available data (e.g., for the Sierra Nevada Scots pine, *Pinus sylvestris*, population in southern Spain, an isolated marginal population at the species southern distribution limit; Alía et al., 2001; Castro et al., 2004). Thus, the potential for both genetic adaptation and plastic responses needs to be integrated in models predicting the responses of marginal populations to climate change. Furthermore, species’ range limits are not determined solely by climate and demographic processes. Loehle (1998) showed that the range limits of many low-latitude tree species are set by competitive interactions with other tree species. Other biotic factors, such as the positive interactions between species (Stephan et al., 2021), are known to have a strong influence on the definition of tree range boundaries. To better disentangle the relationship between genetic indicators and population marginality within a species’ range, future research should address the development of predictive models that include species-specific indicators related to biotic interactions.

## Conclusion

In maritime pine, a trend of decreasing in both overall and climate-associated outlier genetic diversity, and increasing differentiation with geographic marginality supported the well-established centre-periphery hypothesis, at both range-wide and regional (gene pool) spatial scales. However, geographically marginal populations displayed also lower recessive genetic load compared to core populations, which, together with expected novel adaptations in the species range margins, highlight their importance in the context of future adaptation to climate change. Higher genomic offset in ecologically marginal populations suggests higher potential maladaptation of these populations to future climates and the urgency to develop specific management actions to preserve populations in marginal climatic conditions. Overall, our study shows the importance of combining quantitative marginality indices and diverse genetic indicators, gauging for multiple evolutionary processes, to have a sound basis for conservation decisions.

## Author Contribution

SCG-M, MW, GGV, DG, AP and RA conceptualized the study. SP, FB, GGV, CA and SCG-M produced, cleaned and formatted the genetic data. SCG-M, MW, DG and RA conceived the paper methodology. AT, CG-P, JA and SCG-M analysed the data. AT, SCG-M and MW drafted the article. All authors contributed to revision and editing of the final version, and gave their final approval for publication.

## Acknowledgements

We thank Carmen García-Barriga, Francisco Auñón, Eduardo Ballestero, Diana Barba, Maria Regina Chambel, Fernando Del-Caño and Rodrigo Pulido-Sanz for their contribution to genetic data production. We are also very grateful to Dr Bruno Fady and Dr Nicolas Picard for their valuable recommendations and help at different stages of this study.

## Funding statement

This study was funded by the European Union’s Horizon 2020 research and innovation program under grant agreement No 862221 (FORGENIUS). Views and opinions expressed are however those of the author(s) only and do not necessarily reflect those of the European Union. Neither the European Union nor the granting authority can be held responsible for them. The study also received financial support from the French government in the framework of the IdEx Bordeaux University ‘Investments for the Future’ program / GPR Bordeaux Plant Sciences (France) and the MITECO contract for Conservation of Genetic Resources (Spain).

## Data archiving statement

Data are openly available in a public repository (Data INRAE) and was provided for the anonymized review process as a file. Scripts for preparation of data, selection of climate variables and computation of genomic offset statistics can be found in a GitHub public repository, and were cloned and anonymised for the review process at https://anonymous.4open.science/r/ReadyToGO_Pinpin-FA56/README.md.

## Conflict of interest statement

The authors declare that they have no known competing financial interests or personal relationships that could have appeared to influence the work reported in this paper.

## Biosketch

This study is the result of a long-term collaboration among four public research institutions in Spain, Italy, Slovenia and France, which are working together on elucidating the distribution of genetic diversity and patterns of local adaptation in European forest trees, in particular those from highly diverse Mediterranean and Alpine forests. Our research interests are broad, including population and quantitative genetics, biogeography, conservation genetics, ecological modelling and genomics. The paper constitutes the first PhD manuscript of the first author, Adélaïde Theraroz, developed at the University of Bordeaux (France), and supervised by Marjana Westergren (Slovenian Forestry Institute) and Santiago C. González-Martínez (INRAE).

## Short title

Genetic effects of population marginality

## Notes

https://entrepot.recherche.data.gouv.fr/dataset.xhtml?persistentId=doi:10.57745/SPUDQQ

### Competing Interest Statement

The authors have declared no competing interest.

### Summary of Updates

The new version of the manuscript contains additional analyses and results, which have been discussed in the dedicated sections. These changes were requested by the referees of the journal to which we submitted the paper.

## References

Abad-Viñas, R., Caudullo, G., Oliveira, S., & de Rigo, D. (2016). *Pinus pinaster* in Europe: distribution, habitat, usage and threats. In: San-Miguel-Ayanz, J., de Rigo, D., Caudullo, G., Houston Durrant, T., Mauri, A. (Eds.), European Atlas of Forest Tree Species. Publ. Off. EU, Luxembourg, pp. e012d59+.

Ahrens, C. W., Rymer, P. D., & Miller, A. D. (2023). Genetic offset and vulnerability modelling: misinterpretations of results and violations of evolutionary principles. Authorea. 10.22541/au.168727971.18670759/v1.

Alía, R., Martín, S., de Miguel, J., Galera, R., Agúndez, D., Gordo, J., Salvador, L., Catalán, G., & Gil, L. (1996). Las regiones de procedencia de Pinus pinaster Aiton en España (Organismo Autónomo Parques Nacionales-ETSI de Montes).

Alía, R., Moro-Serrano, J., & Notivol, E. (2001). Genetic variability of Scots pine (*Pinus sylvestris*) provenances in Spain: growth traits and survival. Silva Fennica, 35(1), 601. 10.14214/sf.601.

Archambeau, J., Benito-Garzón, M., de-Miguel, M., Changenet, A., Bagnoli, F., Barraquand, F., Marchi, M., Vendramin, G.G., Cavers, S., Perry, A., & González-Martínez, S.G. (2024). Evaluating genomic offset predictions in a forest tree with high population genetic structure. bioRxiv. 10.1101/2024.05.17.594631.

Baradat, P., Marpeau-Bezard, A. (1988). Le pin maritime *Pinus pinaster* Ait. Biologie et génétique des terpènes pour la connaissance et l’amélioration de l’espèce. Thèse doctorale d’État, Université Bordeaux I.

Birch, L. C. (1957). The role of weather in determining the distribution and abundance of animals. Cold Spring Harbor Symposia on Quantitative Biology, 22, 203-218. doi:10.1101/SQB.1957.022.01.021.

Brown, J. H. (1984). On the relationship between abundance and distribution of species. The American Naturalist, 124(2), 255-279. 10.1086/284267.

Bucci, G., González-Martínez, S. C., Le Provost, G., Plomion, C., Ribeiro, M. M., Sebastiani, F., Alía, R., & Vendramin, G. G. (2007). Range-wide phylogeography and gene zones in *Pinus pinaster* Ait. revealed by chloroplast microsatellite markers. Molecular Ecology, 16(10), 2137-2153. 10.1111/j.1365-294X.2007.03275.x.

Burban, C., & Petit, R. J. (2003). Phylogeography of maritime pine inferred with organelle markers having contrasted inheritance. Molecular Ecology, 12(6), 1487-1495. 10.1046/j.1365-294X.2003.01817.x.

Caballero, A., Bravo, I., & Wang, J. (2017). Inbreeding load and purging: implications for the short-term survival and the conservation management of small populations. Heredity, 118(2), 177-185. 10.1038/hdy.2016.80.

Cahn, L. (2023). Genetic basis of local adaptation of maritime pine in Corsica [Unpublished master’s thesis]. Université de Rennes 1.

Capblancq, T., Fitzpatrick, M. C., Bay, R. A., Expósito-Alonso, M., & Keller, S. R. (2020). Genomic prediction of (mal)adaptation across current and future climatic landscapes. Annual Review of Ecology, Evolution, and Systematics, 51(1), 245-269. 10.1146/annurev-ecolsys-020720-042553.

Castro, J., Zamora, R., Hódar, J. A., & Gómez, J. M. (2004). Seedling establishment of a boreal tree species (*Pinus sylvestris*) at its southernmost distribution limit: consequences of being in a marginal Mediterranean habitat. Journal of Ecology, 92(2), 266-277. 10.1111/j.0022-0477.2004.00870.x.

Caudullo, G., Welk, E., & San-Miguel-Ayanz, J. (2017). Chorological maps for the main European woody species. Data in Brief, 12, 662-666. 10.1016/j.dib.2017.05.007.

Cingolani, P., Platts, A., Wang, L. L., Coon, M., Nguyen, T., Wang, L., Land, S. J., Lu, X., & Ruden, D. M. (2012). A program for annotating and predicting the effects of single nucleotide polymorphisms, SnpEff. Fly, 6(2), 80–92. 10.4161/fly.19695.

De La Torre, A. R., Wang, T., Jaquish, B., & Aitken, S. N. (2014). Adaptation and exogenous selection in a *Picea glauca* × *Picea engelmannii* hybrid zone: implications for forest management under climate change. New Phytologist, 201(2), 687-699. 10.1111/nph.12540.

De-Lucas, A. I., González-Martínez, S. C., Vendramin, G. G., Hidalgo, E., & Heuertz, M. (2009). Spatial genetic structure in continuous and fragmented populations of *Pinus pinaster* Aiton. Molecular Ecology, 18(22), 4564-4576. 10.1111/j.1365-294X.2009.04372.x.

De Kort, H., Prunier, J. G., Ducatez, S., Honnay, O., Baguette, M., Stevens, V. M., & Blanchet, S. (2021). Life history, climate and biogeography interactively affect worldwide genetic diversity of plant and animal populations. Nature Communications, 12(1). 10.1038/s41467-021-20958-2.

Dussex, N., Morales, H. E., Grossen, C., Dalén, L., & van Oosterhout, C. (2023). Purging and accumulation of genetic load in conservation. Trends in Ecology & Evolution, 38(10), 961-969. 10.1016/j.tree.2023.05.008.

Dussex, N., van der Valk, T., Morales, H. E., Wheat, C. W., Díez-del-Molino, D., von Seth, J., Foster, Y., Kutschera, V. E., Guschanski, K., Rhie, A., Phillippy, A. M., Korlach, J., Howe, K., Chow, W., Pelan, S., Mendes Damas, J. D., Lewin, H. A., Hastie, A. R., Formenti, G., … Dalén, L. (2021). Population genomics of the critically endangered kākāpō. Cell Genomics, 1(1), 100002. 10.1016/j.xgen.2021.100002.

Eckert, C. G., Samis, K. E., & Lougheed, S. C. (2008). Genetic variation across species’ geographical ranges: the central–marginal hypothesis and beyond. Molecular Ecology, 17(5), 1170-1188. 10.1111/j.1365-294X.2007.03659.x.

Ellis, N., Smith, S. J., & Pitcher, C. R. (2012). Gradient forests: calculating importance gradients on physical predictors. Ecology, 93(1), 156-168. 10.1890/11-0252.1.

Evanno, G., Regnaut, S., & Goudet, J. (2005). Detecting the number of clusters of individuals using the software STRUCTURE: a simulation study. Molecular Ecology, 14(8), 2611-2620. 10.1111/j.1365-294X.2005.02553.x.

Excoffier, L., Foll, M., & Petit, R. J. (2009). Genetic consequences of range expansions. Annual Review of Ecology, Evolution, and Systematics, 40(1), 481–501. 10.1146/annurev.ecolsys.39.110707.173414.

Fady, B., & Conord, C. (2010). Macroecological patterns of species and genetic diversity in vascular plants of the Mediterranean basin. Diversity and Distributions, 16(1), 53–64. 10.1111/j.1472-4642.2009.00621.x.

Feng, S., Fang, Q., Barnett, R., Li, C., Han, S., Kuhlwilm, M., Zhou, L., Pan, H., Deng, Y., Chen, G., Gamauf, A., Woog, F., Prys-Jones, R., Marques-Bonet, T., Gilbert, M. T. P., & Zhang, G. (2019). The genomic footprints of the fall and recovery of the crested Ibis. Current Biology, 29(2), 340–349.e7. 10.1016/j.cub.2018.12.008.

Fitzpatrick, M. C., Chhatre, V. E., Soolanayakanahally, R. Y., & Keller, S. R. (2021). Experimental support for genomic prediction of climate maladaptation using the machine learning approach Gradient Forests. Molecular Ecology Resources, 21(8), 2749-2765. 10.1111/1755-0998.13374.

Fitzpatrick, M. C., & Keller, S. R. (2015). Ecological genomics meets community-level modelling of biodiversity: mapping the genomic landscape of current and future environmental adaptation. Ecology Letters, 18(1), 1-16. 10.1111/ele.12376.

Fkiri, S., Guibal, F., Elkhorchani, A., Khouja, M. L., Khaldi, A., & Nasr, Z. (2019). Relationship between climate and growth of two North African varieties of *Pinus pinaster* Arn. African Journal of Ecology, 57(3), 327-334. 10.1111/aje.12610.

Foll, M., & Gaggiotti, O. (2008). A genome-scan method to identify selected loci appropriate for both dominant and codominant markers: a Bayesian perspective. Genetics, 180(2), 977-993. 10.1534/genetics.108.092221.

Fréjaville, T., Vizcaíno-Palomar, N., Fady, B., Kremer, A., & Benito Garzón, M. (2019). Range margin populations show high climate adaptation lags in European trees. Global Change Biology, 26(2), 484–495. 10.1111/gcb.14881.

Frichot, E., Schoville, S. D., Bouchard, G., & François, O. (2013). Testing for associations between loci and environmental gradients using Latent Factor Mixed Models. Molecular Biology and Evolution, 30(7), 1687-1699. 10.1093/molbev/mst063.

Gamba, D., & Muchhala, N. (2020). Global patterns of population genetic differentiation in seed plants. Molecular Ecology, 29(18), 3413–3428. 10.1111/mec.15575.

Gapare, W. J., Aitken, S. N., & Ritland, C. E. (2005). Genetic diversity of core and peripheral Sitka spruce (*Picea sitchensis* (Bong.) Carr) populations: implications for conservation of widespread species. Biological Conservation, 123(1), 113-123. 10.1016/j.biocon.2004.11.002.

Gaston, K. J. (2003). The structure and dynamics of geographic ranges. Oxford University Press, Oxford.

Gaston, K. J. (2009). Geographic range limits: achieving synthesis. Proceedings of the Royal Society B: Biological Sciences, 276(1661), 1395-1406. 10.1098/rspb.2008.1480.

Gautier, M. (2015). Genome-wide scan for adaptive divergence and association with population-specific covariates. Genetics, 201(4), 1555-1579. 10.1534/genetics.115.181453.

González-Martínez, S.C., Gómez, A., Carrión, J.S., Agúndez, D., Alía, R., Gil, L. (2007). Spatial genetic structure of an explicit glacial refugium of maritime pine (*Pinus pinaster* Aiton) in southeastern Spain. In: Weiss, S., Ferrand, N. (eds) Phylogeography of Southern European Refugia. Springer, Dordrecht.

González-Martínez, S. C., Ridout, K., & Pannell, J. R. (2017). Range expansion compromises adaptive evolution in an outcrossing plant. Current Biology, 27(16), 2544–2551.e4. 10.1016/j.cub.2017.07.007.

Gougherty, A. V., Keller, S. R., & Fitzpatrick, M. C. (2021). Maladaptation, migration and extirpation fuel climate change risk in a forest tree species. Nature Climate Change, 11(2), 166-171. 10.1038/s41558-020-00968-6.

Grossen, C., Guillaume, F., Keller, L. F., & Croll, D. (2020). Purging of highly deleterious mutations through severe bottlenecks in Alpine ibex. Nature Communications, 11(1), 1001. 10.1038/s41467-020-14803-1.

Gugerli, F., Rüegg, M., & Vendramin, G. G. (2009). Gradual decline in genetic diversity in Swiss stone pine populations (*Pinus cembra*) across Switzerland suggests postglacial re-colonization into the Alps from a common eastern glacial refugium. Botanica Helvetica, 119(1), 13-22. 10.1007/s00035-009-0052-6.

Günther, T., & Schmid, K. J. (2010). Deleterious amino acid polymorphisms in *Arabidopsis thaliana* and rice. Theoretical and Applied Genetics, 121(1), 157-168. 10.1007/s00122-010-1299-4.

Hampe, A., & Petit, R. J. (2005). Conserving biodiversity under climate change: the rear edge matters. Ecology Letters, 8(5), 461-467. 10.1111/j.1461-0248.2005.00739.x.

Hedrick, P. W., & García-Dorado, A. (2016). Understanding inbreeding depression, purging, and genetic rescue. Trends in Ecology & Evolution, 31(12), 940-952. 10.1016/j.tree.2016.09.005.

Hengeveld, R., & Haeck, J. (1982). The distribution of abundance. I. Measurements. Journal of Biogeography, 9(4), 303. 10.2307/2844717.

Hilfiker, K., Gugerli, F., Schütz, J.-P., Rotach, P., & Holderegger, R. (2004). Low RAPD variation and female-biased sex ratio indicate genetic drift in small populations of the dioecious conifer *Taxus baccata* in Switzerland. Conservation Genetics, 5(3), 357-365. 10.1023/B:COGE.0000031144.95293.1b.

Hoban, S., Kelley, J. L., Lotterhos, K. E., Antolin, M. F., Bradburd, G., Lowry, D. B., Poss, M. L., Reed, L. K., Storfer, A., & Whitlock, M. C. (2016). Finding the genomic basis of local adaptation: pitfalls, practical solutions, and future directions. The American Naturalist, 188(4), 379–397. 10.1086/688018.

IPCC (2021). Climate Change 2021: The Physical Science Basis: Contribution of Working Group I to the Sixth Assessment Report of the Intergovernmental Panel on Climate Change. Cambridge University Press, Cambridge, United Kingdom and New York, NY, USA.

Jaramillo-Correa, J. P., Grivet, D., Terrab, A., Kurt, Y., De-Lucas, A. I., Wahid, N., Vendramin, G. G., & González-Martínez, S. C. (2010). The Strait of Gibraltar as a major biogeographic barrier in Mediterranean conifers: a comparative phylogeographic survey. Molecular Ecology, 19(24), 5452–5468. 10.1111/j.1365-294x.2010.04912.x.

Jaramillo-Correa, J.-P., Rodríguez-Quilón, I., Grivet, D., Lepoittevin, C., Sebastiani, F., Heuertz, M., Garnier-Géré, P. H., Alía, R., Plomion, C., Vendramin, G. G., & González-Martínez, S. C. (2015). Molecular proxies for climate maladaptation in a long-lived tree (*Pinus pinaster* Aiton, Pinaceae). Genetics, 199(3), 793-807. 10.1534/genetics.114.173252.

Jia, K., Zhao, W., Maier, P. A., Hu, X., Jin, Y., Zhou, S., Jiao, S., El-Kassaby, Y. A., Wang, T., Wang, X., & Mao, J. (2020). Landscape genomics predicts climate change-related genetic offset for the widespread *Platycladus orientalis* (Cupressaceae). Evolutionary Applications, 13(4), 665-676. 10.1111/eva.12891.

Johannesson, K., & André, C. (2006). Life on the margin: genetic isolation and diversity loss in a peripheral marine ecosystem, the Baltic Sea. Molecular Ecology, 15(8), 2013-2029. 10.1111/j.1365-294X.2006.02919.x.

Jost, L. (2008). *G*_ST_ and its relatives do not measure differentiation. Molecular Ecology, 17(18), 4015-4026. 10.1111/j.1365-294X.2008.03887.x.

Kimura, M., Maruyama, T., & Crow, J. F. (1963). The mutation load in small populations. Genetics, 48(10), 1303–1312. 10.1093/genetics/48.10.1303.

Kimura, M., & Ohta, T. (1969). The average number of generations until fixation of a mutant gene in a finite population. Genetics, 61(3), 763-771. 10.1093/genetics/61.3.763.

Kolzenburg, R. (2022). The direct influence of climate change on marginal populations: a review. Aquatic Sciences, 84(2), 24. 10.1007/s00027-022-00856-5.

Lande, R. (1988). Genetics and demography in biological conservation. Science, 241(4872), 1455-1460. 10.1126/science.3420403.

Láruson, Á. J., Fitzpatrick, M. C., Keller, S. R., Haller, B. C., & Lotterhos, K. E. (2022). Seeing the forest for the trees: Assessing genetic offset predictions from gradient forest. Evolutionary Applications, 15, 403–416. 10.1111/eva.13354.

Latta, R. G. (1995). The effects of embryo competition with mixed mating on the genetic load in plants. Heredity, 75(6), 637-643. 10.1038/hdy.1995.183.

Lind, B. M., Candido-Ribeiro, R., Singh, P., Lu, M., Vidaković, D. O., Booker, T. R., Whitlock, M. C., Yeaman, S., Isabel, N., & Aitken, S. N. (2024). How useful is genomic data for predicting maladaptation to future climate? Global Change Biology, 30(4). 10.1111/gcb.17227.

Lira-Noriega, A., & Manthey, J. D. (2014). Relationship of genetic diversity and niche centrality: a survey and analysis. Evolution, 68(4), 1082-1093. 10.1111/evo.12343.

Loehle, C. (1998). Height growth rate trade-offs determine northern and southern range limits for trees. Journal of Biogeography, 25(4), 735–742. http://www.jstor.org/stable/2846146.

Lotterhos, K. E., & Whitlock, M. C. (2014). Evaluation of demographic history and neutral parameterization on the performance of *F*_ST_ outlier tests. Molecular Ecology, 23(9), 2178–2192. 10.1111/mec.12725.

Lotterhos, K. E., & Whitlock, M. C. (2015). The relative power of genome scans to detect local adaptation depends on sampling design and statistical method. Molecular Ecology, 24(5), 1031–1046. 10.1111/mec.13100.

Lotterhos, K. E. and Whitlock, M. C. (2014). Evaluation of demographic history and neutral parameterization on the performance of FST outlier tests. Molecular Ecology, 23, 2178– 2192. 10.1111/mec.12725.

Lotterhos, K. E. (2024). Interpretation issues with “genomic vulnerability” arise from conceptual issues in local adaptation and maladaptation. Evolution Letters, 8(3), 331–339. 10.1093/evlett/qsrae004.

Lynch, M., Conery, J., & Bürger, R. (1995). Mutation accumulation and the extinction of small populations. The American Naturalist, 146(4), 489-518. 10.1086/285812.

Marchi, M., Castellanos-Acuña, D., Hamann, A., Wang, T., Ray, D., & Menzel, A. (2020). ClimateEU, scale-free climate normals, historical time series, and future projections for Europe. Scientific Data, 7(1), 428. 10.1038/s41597-020-00763-0.

Marques, H., Pinto, G., Pinto, P., Teixeira, C. (2012). Regiões de Proveniência. Portugal. ICNF, 47-51.

Milesi, P., Kastally, C., Dauphin, B., Cervantes, S., Bagnoli, F., Budde, K. B., Cavers, S., Fady, B., Faivre-Rampant, P., González-Martínez, S. C., Grivet, D., Gugerli, F., Jorge, V., Kupin, I. L., Ojeda, D. I., Olsson, S., Opgenoorth, L., Pinosio, S., Plomion, C., … Pyhäjärvi, T. (2023). Synchronous effective population size changes and genetic stability of forest trees through glacial cycles. bioRxiv. 10.1101/2023.01.05.522822.

Naydenov, K. D., Alexandrov, A., Matevski, V., Vasilevski, K., Naydenov, M. K., Gyuleva, V., Carcaillet, C., Wahid, N. and Kamary, S. (2014). Range-wide genetic structure of maritime pine predates the last glacial maximum: evidence from nuclear DNA. Hereditas, 151(1), 1–13. 10.1111/j.1601-5223.2013.00027.x.

Nicholson, A. J. (1958). Dynamics of insect populations. Annual Review of Entomology, 3(1), 107-136. 10.1146/annurev.en.03.010158.000543.

Peischl, S., Dupanloup, I., Kirkpatrick, M., & Excoffier, L. (2013). On the accumulation of deleterious mutations during range expansions. Molecular Ecology, 22(24), 5972-5982. 10.1111/mec.12524.

Picard, N., Marchi, M., Serra-Varela, M. J., Westergren, M., Cavers, S., Notivol, E., Piotti, A., Alizoti, P., Bozzano, M., González-Martínez, S. C., Grivet, D., Aravanopoulos, F. A., Vendramin, G. G., Ducci, F., Fady, B., & Alía, R. (2022). Marginality indices for biodiversity conservation in forest trees. Ecological Indicators, 143, 109367. 10.1016/j.ecolind.2022.109367.

Pironon, S., Villellas, J., Morris, W. F., Doak, D. F., & García, M. B. V. (2015). Do geographic, climatic or historical ranges differentiate the performance of central versus peripheral populations? Global Ecology and Biogeography, 24(6), 611-620. 10.1111/geb.12263.

Pironon, S., Papuga, G., Villellas, J., Angert, A. L., García, M. B. V., & Thompson, J. D. (2016). Geographic variation in genetic and demographic performance: new insights from an old biogeographical paradigm. Biological Reviews, 92(4), 1877-1909. 10.1111/brv.12313.

Plomion, C., Bartholomé, J., Lesur, I., Boury, C., Rodríguez-Quilón, I., Lagraulet, H., & González-Martínez, S. C. (2016). High-density SNP assay development for genetic analysis in maritime pine (*Pinus pinaster*). Molecular Ecology Resources, 16(2), 574–587. 10.1111/1755-0998.12464.

Pritchard, J. K., Stephens, M., & Donnelly, P. (2000). Inference of population structure using multilocus genotype data. Genetics, 155(2), 945-959. 10.1093/genetics/155.2.945.

R Core Team. (2022). R: A Language and Environment for Statistical Computing. R Foundation for Statistical Computing. https://www.R-project.org/.

Rellstab, C., Dauphin, B., & Expósito-Alonso, M. (2021). Prospects and limitations of genomic offset in conservation management. Evolutionary Applications, 14(5), 1202-1212. 10.1111/eva.13205.

Richards, O. W. (1961). The theoretical and practical study of natural insect populations. Annual Review of Entomology, 6(1), 147-162. 10.1146/annurev.en.06.010161.001051.

Robinson, J. A., Brown, C., Kim, B. Y., Lohmueller, K. E., & Wayne, R. K. (2018). Purging of strongly deleterious mutations explains long-term persistence and absence of inbreeding depression in island foxes. Current Biology, 28(21), 3487–3494.e4. 10.1016/j.cub.2018.08.066.

Rousset, F. (2008). Genepop’007: a complete re-implementation of the genepop software for Windows and Linux. Molecular Ecology Resources, 8(1), 103-106. 10.1111/j.1471-8286.2007.01931.x.

Sachdeva, H., Olusanya, O., & Barton, N. (2022). Genetic load and extinction in peripheral populations: the roles of migration, drift and demographic stochasticity. Philosophical Transactions of the Royal Society B: Biological Sciences, 377(1846). 10.1098/rstb.2021.0010.

Salvador, L., Alía, R., Agúndez, D., & Gil, L. (2000). Genetic variation and migration pathways of maritime pine (*Pinus pinaster* Ait) in the Iberian Peninsula. Theoretical and Applied Genetics, 100(1), 89–95. 10.1007/s001220050013.

Santos, A. S., & Gaiotto, F. A. (2020). Knowledge status and sampling strategies to maximize cost-benefit ratio of studies in landscape genomics of wild plants. Scientific Reports, 10(1), 3706. 10.1038/s41598-020-60788-8.

Sexton, J. P., McIntyre, P. J., Angert, A. L., & Rice, K. J. (2009). Evolution and ecology of species range limits. Annual Review of Ecology, Evolution, and Systematics, 40(1), 415-436. 10.1146/annurev.ecolsys.110308.120317.

Sexton, J. P., Hangartner, S. B., & Hoffmann, A. A. (2013). Genetic isolation by environment or distance: which pattern of gene flow is most common? Evolution, 68(1), 1-15. 10.1111/evo.12258.

Sgrò, C. M., Lowe, A. J., & Hoffmann, A. A. (2010). Building evolutionary resilience for conserving biodiversity under climate change. Evolutionary Applications, 4(*2*), 326–337. 10.1111/j.1752-4571.2010.00157.x.

Soille, P., & Vogt, P. (2009). Morphological segmentation of binary patterns. Pattern Recognition Letters, 30(4), 456-459. 10.1016/j.patrec.2008.10.015.

Soulé, M.E. (1973). The epistasis cycle: a theory of marginal populations. Annual Review of Ecology and Systematics, 4(1), 165-187. 10.1146/annurev.es.04.110173.001121.

Stephan, P., Mora, B. B., & Alexander, J. M. (2021). Positive species interactions shape species’ range limits. Oikos, 130(10), 1611–1625. 10.1111/oik.08146.

Travis, J. M. J., Munkemuller, T., Burton, O. J., Best, A., Dytham, C., & Johst, K. (2007). Deleterious mutations can surf to high densities on the wave front of an expanding population. Molecular Biology and Evolution, 24(10), 2334–2343. 10.1093/molbev/msm167.

van den Wollenberg, A. L. (1977). Redundancy analysis an alternative for canonical correlation analysis. Psychometrika, 42(2), 207-219. 10.1007/BF02294050.

Vendramin, G., Anzidei, M., Madaghiele, A., Bucci, G. (1998). Distribution of genetic diversity in *Pinus pinaster* Ait. as revealed by chloroplast microsatellites. Theoretical and Applied Genetics, 97, 456–463. 10.1007/s001220050917.

Wahid, N., González-Martínez, S. C., Hadrami, I. E., & Boulli, A. (2004). Genetic Structure and variability of natural populations of maritime pine (*Pinus pinaster* Aiton) in Morocco. Silvae Genetica, *53*(1-6), 93-99. 10.1515/sg-2004-0017.

Wahid, N., González-Martínez, S. C., El Hadrami, I., & Boulli, A. (2006). Variation of morphological traits in natural populations of maritime pine (*Pinus pinaster* Ait.) in Morocco. Annals of Forest Science, 63(1), 83-92. 10.1051/forest:20050100.

Weir, B. S., & Cockerham, C. C. (1984). Estimating F-statistics for the analysis of population structure. Evolution, 38(6), 1358. 10.2307/2408641.

Westergren, M., Božič, G., Kraigher, H. (2018). Genetic diversity of core *vs.* peripheral Norway spruce native populations at a local scale in Slovenia. iForest - Biogeosciences and Forestry, 11(1), 104–110. 10.3832/ifor2444-011.

Whittaker, J. B. (1971). Population changes in *Neophilaenus lineatus* (L.) (Homoptera : Cercopidae) in different parts of its range. The Journal of Animal Ecology, 40(2), 425. 10.2307/3253.

Willson, M. F., & Burley, N. (1983). *Mate Choice in Plants (MPB-19), Volume 19: Tactics, Mechanisms, and Consequences. (MPB-19)*. Princeton University Press. 10.2307/j.ctvx5wbss.

Young, A.G., Boyle, T., & Brown, T.L. (1996). The population genetic consequences of habitat fragmentation for plants. Trends in Ecology & Evolution, 11(10), 413-418. 10.1016/0169-5347(96)10045-8.

Yuan, S., Shi, Y., Zhou, B., Liang, Y., Chen, X., An, Q., Fan, Y., Shen, Z., Ingvarsson, P. K., & Wang, B. (2023). Genomic vulnerability to climate change in *Quercus acutissima*, a dominant tree species in East Asian deciduous forests. Molecular Ecology, 32(7), 1639-1655. 10.1111/mec.16843.

