## Supplementary Files for "The genetic consequences of population marginality: a case study in maritime pine"

**Table S1**. Geographical location and sample size (*N*) of the 82 populations of *P. pinaster* included in the study; Abb.: Abbreviated population name.

| Population | Abb. | Country | *N* | Latitude (°) | Longitude (°) | Elevation (m) |
|---|---|---|---|---|---|---|
| Aïn Babouch | ABA | Tunisia | 22 | 36.815 | 8.715 | 201 |
| Adeldal | ADE | Morocco | 21 | 35.157 | -5.079 | 830 |
| Ahin | AHI | Spain | 22 | 39.888 | -0.332 | 733 |
| Alto de la Llama | ALT | Spain | 8 | 43.297 | -6.463 | 526 |
| Ania | ANI | France | 12 | 41.968 | 9.284 | 578 |
| Arenzano | ANO | Italy | 11 | 44.418 | 8.671 | 403 |
| Armayán | ARM | Spain | 8 | 43.305 | -6.458 | 559 |
| Arenas de San Pedro | ARN | Spain | 17 | 40.195 | -5.116 | 663 |
| Bavella | BAV | France | 16 | 41.796 | 9.235 | 1042 |
| Bayubas de Abajo | BAY | Spain | 15 | 41.523 | -2.877 | 925 |
| Benicassim | BEN | Spain | 22 | 40.079 | 0.025 | 520 |
| Bonifatu | BOI | France | 16 | 42.446 | 8.825 | 500 |
| Boniches | BON | Spain | 8 | 39.984 | -1.662 | 1090 |
| Cadavedo | CAD | Spain | 8 | 43.540 | -6.418 | 164 |
| Cagna | CAG | France | 12 | 41.597 | 9.142 | 995 |
| Carbonero el Mayor | CAR | Spain | 6 | 41.172 | -4.277 | 844 |
| Castropol | CAS | Spain | 8 | 43.501 | -6.983 | 158 |
| Cazorla | CAZ | Spain | 21 | 37.918 | -2.927 | 1059 |
| Cenicientos | CEN | Spain | 9 | 40.278 | -4.491 | 1079 |
| Coca | COC | Spain | 17 | 41.255 | -4.498 | 784 |
| Codos | COD | Spain | 22 | 41.282 | -1.418 | 1058 |
| Cómpeta | COM | Spain | 3 | 36.832 | -3.925 | 904 |

| | | | | | | |
|---|---|---|---|---|---|---|
| Cortes de Pallás | CPA | Spain | 22 | 39.178 | -0.946 | 925 |
| Cuéllar | CUE | Spain | 23 | 41.335 | -4.249 | 826 |
| Estepona | EST | Spain | 21 | 36.516 | -5.121 | 458 |
| Fuencaliente | FUE, stand 1 | Spain | 21 | 38.417 | -4.254 | 913 |
| | FUE, stand 2 | Spain | 25 | 38.417 | -4.254 | 913 |
| Gaucin | GAU | Spain | 22 | 36.532 | -5.301 | 630 |
| Gea de Albarracín | GEA | Spain | 22 | 40.365 | -1.351 | 1367 |
| La Bisbal | GIR | Spain | 21 | 41.899 | 3.032 | 243 |
| Guagno | GUA | France | 12 | 42.173 | 8.879 | 572 |
| Hourtin | HOU | France | 25 | 45.183 | -1.150 | 28 |
| Jubrique | JUB | Spain | 21 | 36.523 | -5.185 | 958 |
| Koudiat Erramla | KUD | Morocco | 16 | 35.467 | -5.383 | 480 |
| F.D. de Lacanau | LAC | France | 21 | 44.946 | -1.187 | 36 |
| Lamuño | LAM | Spain | 9 | 43.559 | -6.219 | 119 |
| Les Corbières | LCO, stand 1 | France | 21 | 43.082 | 2.878 | 216 |
| | LCO, stand 2 | France | 25 | 43.082 | 2.878 | 216 |
| Leiria | LEI | Portugal | 21 | 39.783 | -8.958 | 79 |
| F.D. de Lit et Mixe | LIM | France | 25 | 44.052 | -1.301 | 45 |
| Madisouka | MAD, stand 1 | Morocco | 1 | 35.191 | -5.166 | 1302 |
| | MAD, stand 2 | Morocco | 21 | 35.191 | -5.166 | 1302 |
| Magra/Montemarcello | MAG | Italy | 15 | 44.074 | 9.973 | 100 |
| Maures | MAU, stand 1 | France | 7 | 43.233 | 6.365 | 361 |
| | MAU, stand 2 | France | 8 | 43.233 | 6.365 | 606 |
| Mazarete | MAZ | Spain | 21 | 40.972 | -2.222 | 1171 |
| Mimizan | MIM | France | 18 | 44.134 | -1.303 | 18 |

| | | | | | | |
|---|---|---|---|---|---|---|
| Montignoso | MON | Italy | 6 | 44.012 | 10.184 | 434 |
| Pina de Montalgrao | MTG | Spain | 21 | 40.030 | -0.644 | 1148 |
| Murlo | MUR | Italy | 8 | 43.129 | 11.222 | 408 |
| Olba | OLB | Spain | 20 | 40.173 | -0.623 | 989 |
| Olonne sur Mer | OLO | France | 24 | 46.566 | -1.831 | 12 |
| Oña | ONA | Spain | 24 | 42.763 | -3.535 | 753 |
| Oria | ORI | Spain | 23 | 37.513 | -2.317 | 1221 |
| Pantelleria | PAN | Italy | 17 | 36.782 | 12.002 | 733 |
| Point Cires | PCI | Morocco | 22 | 35.904 | -5.481 | 90 |
| Pradell de la Teixeta | PDL | Spain | 22 | 41.166 | 0.865 | 560 |
| Petrocq | PET | France | 22 | 44.064 | -1.300 | 21 |
| La Peza | PEZ | Spain | 22 | 37.263 | -3.403 | 1414 |
| Pinofranqueado | PFQ | Spain | 21 | 40.365 | -6.385 | 724 |
| Pineto | PIN | France | 12 | 42.427 | 9.227 | 475 |
| Pleucadec | PLE | France | 19 | 47.781 | -2.344 | 70 |
| Puerto de Vega | PUE | Spain | 7 | 43.548 | -6.631 | 81 |
| Quatretonda | QUA | Spain | 16 | 38.990 | -0.349 | 420 |
| Quintana Redonda | QUI | Spain | 22 | 41.533 | -2.583 | 1046 |
| Riopar | RIO | Spain | 24 | 38.485 | -2.425 | 1039 |
| Rossiglione | ROS | Italy | 10 | 44.451 | 8.675 | 955 |
| San Cipriano de Ribarteme | SAC | Spain | 10 | 42.116 | -8.366 | 386 |
| El Sahugo | SAH | Spain | 22 | 40.381 | -6.562 | 817 |
| San Leonardo de Yagüe | SAL | Spain | 10 | 41.833 | -3.064 | 1070 |
| Sierra Calderona | SCD | Spain | 22 | 39.749 | -0.495 | 746 |
| Seborga | SEB | Italy | 24 | 43.820 | 7.710 | 544 |

| | | | | | | |
|---|---|---|---|---|---|---|
| Sergude | SEG | Spain | 19 | 42.817 | -8.450 | 309 |
| Sidi-Meskour | SID | Morocco | 22 | 31.506 | -6.995 | 1975 |
| Sierra de Barcia | SIE | Spain | 8 | 43.528 | -6.493 | 264 |
| Sinarcas | SIN | Spain | 22 | 39.791 | -1.203 | 888 |
| Sobron | SOB | Spain | 24 | 42.789 | -3.086 | 895 |
| St-Jean des Monts | STJ | France | 25 | 46.764 | -2.029 | 6 |
| Tamjout | TAJ | Morocco | 22 | 33.842 | -4.007 | 1500 |
| Talayuela | TAL, stand 1 | Spain | 21 | 40.000 | -5.623 | 272 |
| | TAL, stand 2 | Spain | 25 | 40.000 | -5.623 | 272 |
| Tamrabta | TAM | Morocco | 14 | 33.600 | -5.017 | 1729 |
| Tabuyo del Monte | TBY | Spain | 22 | 42.294 | -6.212 | 988 |
| Tocchi | TOC | Italy | 12 | 43.132 | 11.243 | 441 |
| Le Verdon | VER | France | 25 | 45.552 | -1.091 | 10 |
| Villamalur | VMA | Spain | 21 | 39.965 | -0.401 | 639 |
| Valdemaqueda | VMQ | Spain | 10 | 40.514 | -4.313 | 947 |

**Table S2**. Standardized marginality indices for each of the 81 *P. pinaster* populations included in the analysis (population COM was removed due to low sample size), calculated according to Picard et al. (2022) but based on the native range of the species only (i.e., excluding plantations; see Figure S1), and ecological index based on climate distance.

| Population | Area | Centroid | Edge | Gravity | Isolation | Second nearest-core | North-South | East-West | Ecological index |
|---|---|---|---|---|---|---|---|---|---|
| ABA | -0.287 | 0.339 | 1.301 | -0.428 | -0.920 | -0.858 | -0.927 | 0.501 | -0.507 |
| ADE | -0.265 | -1.028 | 0.888 | -0.252 | -0.899 | -0.919 | -0.979 | -1.396 | 1.757 |
| AHI | -0.311 | -1.625 | 1.310 | -0.472 | -1.335 | -1.085 | -0.519 | 0.053 | -1.172 |
| ALT | -0.313 | -1.382 | 0.970 | 0.785 | -0.942 | -1.001 | 0.983 | -1.544 | 0.516 |
| ANI | -0.315 | -0.152 | 1.307 | -0.482 | -1.353 | -0.810 | 0.830 | 0.626 | -0.718 |
| ANO | -0.034 | -0.780 | 1.331 | 0.052 | -1.205 | -1.166 | 1.216 | 0.498 | -0.350 |
| ARM | -0.313 | -1.382 | 0.970 | 0.785 | -0.942 | -1.001 | 0.983 | -1.544 | 0.589 |
| ARN | 0.004 | -1.630 | 1.312 | 0.124 | -1.299 | -1.193 | -0.315 | -1.408 | 0.364 |
| BAV | -0.315 | -0.143 | 1.294 | -0.472 | -1.355 | -0.767 | 0.738 | 0.621 | -0.530 |
| BAY | 0.348 | -1.730 | 1.313 | 0.779 | -1.230 | -1.194 | 0.515 | -0.773 | -0.305 |
| BEN | -0.311 | -1.593 | 1.217 | -0.324 | -1.272 | -1.182 | -0.393 | 0.065 | -1.198 |
| BOI | -0.102 | -0.167 | 1.301 | -0.079 | -1.207 | -1.169 | 0.902 | 0.515 | -0.721 |
| BON | 1.057 | -1.698 | 1.322 | 2.125 | -1.271 | -1.110 | -0.448 | -0.268 | -0.227 |

| | | | | | | | | | |
|---|---|---|---|---|---|---|---|---|---|
| CAD | -0.313 | -1.363 | 0.897 | 0.661 | -0.869 | -0.928 | 1.040 | -1.535 | 1.500 |
| CAG | -0.317 | -0.128 | 1.263 | -0.441 | -1.291 | -0.702 | 0.585 | 0.599 | -1.000 |
| CAR | 1.339 | -1.687 | 1.326 | 2.659 | -1.206 | -1.167 | 0.225 | -1.095 | -0.350 |
| CAS | -0.313 | -1.333 | 0.871 | 0.775 | -0.903 | -0.899 | 1.030 | -1.603 | 1.400 |
| CAZ | 0.246 | -1.527 | 1.316 | 0.584 | -1.171 | -1.130 | -0.875 | -0.787 | 0.113 |
| CEN | 0.004 | -1.652 | 1.301 | 0.125 | -1.235 | -1.199 | -0.249 | -1.185 | 0.289 |
| COC | 1.339 | -1.675 | 1.380 | 2.659 | -1.169 | -1.128 | 0.289 | -1.185 | -0.266 |
| COD | -0.328 | -1.681 | 1.301 | -0.502 | -1.210 | -1.172 | 0.325 | -0.218 | -0.415 |
| CPA | -0.210 | -1.629 | 1.316 | -0.281 | -1.327 | -1.135 | -0.746 | -0.046 | -0.893 |
| CUE | 1.339 | -1.685 | 1.371 | 2.659 | -1.160 | -1.118 | 0.359 | -1.083 | -0.237 |
| EST | -0.265 | -1.275 | 1.316 | -0.385 | -1.060 | -1.010 | -0.971 | -1.408 | 1.526 |
| FUE | 0.246 | -1.438 | 0.966 | 0.001 | -0.978 | -0.994 | -0.810 | -1.083 | 0.468 |
| GAU | -0.265 | -1.266 | 1.291 | -0.381 | -1.019 | -0.965 | -0.971 | -1.447 | 0.465 |
| GEA | -0.296 | -1.678 | 1.309 | -0.446 | -1.271 | -1.149 | -0.172 | -0.206 | -0.343 |
| GIR | 0.881 | -1.268 | 0.516 | -0.037 | -0.459 | -0.540 | 0.796 | 0.068 | -0.823 |
| GUA | -0.102 | -0.164 | 1.301 | -0.086 | -1.276 | -0.786 | 0.870 | 0.533 | -0.591 |
| HOU | -0.319 | -1.325 | 1.192 | -0.440 | -1.255 | -1.169 | 1.242 | -0.117 | -0.422 |
| JUB | -0.265 | -1.271 | 1.312 | -0.385 | -1.044 | -0.992 | -0.971 | -1.426 | 0.515 |

| | | | | | | | | | |
|---|---|---|---|---|---|---|---|---|---|
| KUD | -0.265 | -1.076 | 0.986 | -0.224 | -0.819 | -1.018 | -0.976 | -1.458 | 2.912 |
| LAC | -0.319 | -1.343 | 1.269 | -0.470 | -1.279 | -1.143 | 1.241 | -0.135 | -0.277 |
| LAM | -0.313 | -1.372 | 0.894 | 0.618 | -0.853 | -0.925 | 1.044 | -1.515 | 1.612 |
| LCO | 0.881 | -1.232 | 0.594 | -0.013 | -0.422 | -0.619 | 0.925 | 0.068 | -0.855 |
| LEI | 5.369 | -1.393 | 1.307 | 10.308 | -1.080 | -1.032 | -0.585 | -2.920 | 1.708 |
| LIM | -0.234 | -1.419 | 1.301 | -0.327 | -1.206 | -1.168 | 1.172 | -0.180 | 0.125 |
| MAD | -0.265 | -1.033 | 0.901 | -0.246 | -0.886 | -0.932 | -0.977 | -1.426 | 0.156 |
| MAG | -0.243 | -0.695 | 1.316 | -0.344 | -1.329 | -1.197 | 1.175 | 0.657 | -0.771 |
| MAU | 0.881 | -0.912 | 1.326 | 1.789 | -1.035 | -0.983 | 0.965 | 0.206 | -0.579 |
| MAZ | -0.199 | -1.756 | 1.311 | -0.260 | -1.258 | -1.157 | 0.111 | -0.531 | -0.032 |
| MIM | -0.234 | -1.414 | 1.309 | -0.326 | -1.229 | -1.193 | 1.183 | -0.180 | 0.082 |
| MON | -0.328 | -0.685 | 1.301 | -0.487 | -1.360 | -1.144 | 1.167 | 0.664 | -0.668 |
| MTG | -0.269 | -1.642 | 1.307 | -0.394 | -1.307 | -1.176 | -0.420 | 0.014 | -0.918 |
| MUR | -0.123 | -0.592 | 1.316 | -0.116 | -1.132 | -1.088 | 0.935 | 0.741 | 0.052 |
| OLB | -0.269 | -1.633 | 1.307 | -0.394 | -1.327 | -1.190 | -0.330 | 0.018 | -1.052 |
| OLO | -0.328 | -1.113 | 0.936 | -0.244 | -0.835 | -0.980 | 1.247 | -0.319 | 0.205 |
| ONA | -0.270 | -1.600 | 1.309 | -0.396 | -1.130 | -1.086 | 0.912 | -0.903 | -0.540 |
| ORI | 0.246 | -1.472 | 1.123 | -0.022 | -1.180 | -1.153 | -0.880 | -0.561 | -0.017 |

| | | | | | | | | | |
|---|---|---|---|---|---|---|---|---|---|
| PAN | -0.287 | 0.964 | 0.517 | -0.244 | -0.125 | -0.539 | -0.937 | 0.750 | 1.522 |
| PCI | -0.265 | -1.133 | 1.117 | -0.245 | -0.896 | -1.149 | -0.976 | -1.464 | 5.013 |
| PDL | -0.317 | -1.488 | 0.895 | -0.123 | -0.921 | -0.922 | 0.225 | 0.066 | -0.876 |
| PET | -0.234 | -1.417 | 1.307 | -0.326 | -1.212 | -1.174 | 1.175 | -0.180 | 0.122 |
| PEZ | -0.322 | -1.434 | 1.307 | -0.494 | -1.242 | -1.122 | -0.882 | -0.881 | 0.065 |
| PFQ | -0.181 | -1.533 | 1.318 | -0.225 | -1.266 | -1.171 | -0.172 | -1.531 | -0.334 |
| PIN | -0.102 | -0.186 | 1.307 | -0.075 | -1.238 | -1.197 | 0.899 | 0.621 | -0.495 |
| PLE | -0.328 | -0.957 | 0.537 | -0.115 | -0.457 | -0.575 | 1.248 | -0.576 | -0.107 |
| PUE | -0.313 | -1.349 | 0.882 | 0.689 | -0.871 | -0.911 | 1.044 | -1.559 | 1.588 |
| QUA | -0.210 | -1.582 | 1.202 | 0.024 | -1.200 | -1.161 | -0.761 | 0.048 | -1.325 |
| QUI | 0.348 | -1.724 | 1.313 | 0.779 | -1.239 | -1.127 | 0.515 | -0.676 | -0.314 |
| RIO | 0.246 | -1.573 | 1.328 | 0.584 | -1.269 | -1.004 | -0.803 | -0.605 | 0.313 |
| ROS | -0.034 | -0.778 | 1.320 | 0.052 | -1.202 | -1.164 | 1.222 | 0.498 | -0.255 |
| SAC | 5.369 | -1.410 | 1.288 | 9.639 | -1.286 | -1.120 | 0.863 | -2.581 | 0.068 |
| SAH | -0.181 | -1.525 | 1.307 | -0.225 | -1.302 | -1.193 | -0.159 | -1.551 | -0.509 |
| SAL | 0.348 | -1.707 | 1.326 | 0.779 | -1.274 | -1.075 | 0.762 | -0.828 | -0.175 |
| SCD | -0.327 | -1.631 | 1.301 | -0.481 | -1.349 | -1.140 | -0.608 | 0.026 | -1.308 |
| SEB | -0.034 | -0.836 | 1.309 | 0.052 | -1.251 | -0.910 | 1.118 | 0.434 | -0.745 |

| | | | | | | | | | |
|---|---|---|---|---|---|---|---|---|---|
| SEG | 5.369 | -1.326 | 1.065 | 2.527 | -1.096 | -1.099 | 0.917 | -2.646 | 0.633 |
| SID | -0.323 | -0.513 | 0.565 | -0.255 | -0.644 | -0.586 | -0.996 | -1.603 | -0.630 |
| SIE | -0.313 | -0.513 | 0.565 | -0.255 | -0.644 | -0.586 | -0.996 | -1.603 | 1.363 |
| SIN | 1.057 | -1.674 | 1.343 | 2.125 | -1.248 | -1.190 | -0.585 | -0.135 | -0.539 |
| SOB | -0.270 | -1.599 | 1.258 | -0.267 | -1.125 | -1.080 | 0.916 | -0.834 | -0.345 |
| STJ | -0.328 | -1.089 | 0.861 | -0.208 | -0.768 | -0.904 | 1.247 | -0.402 | 0.024 |
| TAJ | -0.322 | -0.816 | 1.022 | -0.386 | -1.080 | -1.051 | -0.982 | -1.008 | -0.161 |
| TAL | -0.306 | -1.599 | 1.307 | -0.464 | -1.360 | -1.157 | -0.448 | -1.478 | 0.705 |
| TAM | -0.322 | -0.840 | 1.301 | -0.492 | -1.330 | -0.416 | -0.984 | -1.372 | -0.680 |
| TBY | -0.313 | -1.510 | 1.290 | -0.352 | -1.176 | -1.136 | 0.886 | -1.515 | -0.776 |
| TOC | -0.123 | -0.590 | 1.313 | -0.116 | -1.127 | -1.083 | 0.935 | 0.745 | 0.014 |
| VER | -0.328 | -1.297 | 1.301 | -0.505 | -1.153 | -1.110 | 1.245 | -0.094 | -0.652 |
| VMA | -0.311 | -1.630 | 1.301 | -0.442 | -1.361 | -1.112 | -0.461 | 0.042 | -1.114 |
| VMQ | 0.004 | -1.671 | 1.318 | 0.124 | -1.315 | -1.048 | -0.089 | -1.109 | 0.334 |

**Table S3**. Genetic indicators for each of the 81 populations of *P. pinaster* included in the study (population COM was removed due to low sample size). Values for stands within populations were averaged (see Table S1). Overall genetic diversity: genetic diversity estimated as $1\text{-}Q_{inter}$ using all SNPs and corrected for sample size (see main text for details); Genetic diversity for GEA outliers: genetic diversity estimate based on 73 outlier SNPs selected from the GEA performed for the genomic offset computation; Genetic diversity for general outliers: genetic diversity estimate based on 151 outlier SNPs common to two environment-independent outlier-detection methods (see main text for details); $F_{IS}$: population inbreeding; Population-specific $F_{ST}$: genetic differentiation of each population from the common ancestral gene pool; Recessive genetic load: number of predicted deleterious mutations in homozygosity standardized by synonymous mutations; Genomic offset: change in genetic composition required to maintain the current gene-environment relationships under future climates.

| Population | Overall genetic diversity | Genetic diversity for GEA outliers | Genetic diversity for general outliers | $F_{IS}$ | Population-specific $F_{ST}$ | Recessive genetic load | Genomic offset |
|---|---|---|---|---|---|---|---|
| ABA | 0.147 | 0.276 | 0.220 | 0.045 | 0.440 | 0.444 | 0.020 |
| ADE | 0.200 | 0.149 | 0.125 | -0.060 | 0.332 | 0.469 | 0.024 |
| AHI | 0.241 | 0.274 | 0.145 | 0.057 | 0.048 | 0.481 | 0.016 |
| ALT | 0.247 | 0.358 | 0.120 | -0.010 | 0.098 | 0.480 | 0.087 |
| ANI | 0.209 | 0.359 | 0.400 | -0.080 | 0.207 | 0.453 | 0.015 |
| ANO | 0.196 | 0.205 | 0.299 | -0.040 | 0.259 | 0.436 | 0.018 |
| ARM | 0.230 | 0.351 | 0.119 | -0.030 | 0.153 | 0.496 | 0.084 |
| ARN | 0.241 | 0.329 | 0.121 | 0.006 | 0.032 | 0.492 | 0.056 |
| BAV | 0.212 | 0.339 | 0.382 | 0.024 | 0.192 | 0.447 | 0.014 |
| BAY | 0.240 | 0.285 | 0.115 | -0.040 | 0.030 | 0.465 | 0.027 |
| BEN | 0.230 | 0.318 | 0.147 | -0.010 | 0.087 | 0.480 | 0.018 |
| BOI | 0.198 | 0.283 | 0.353 | 0.038 | 0.249 | 0.449 | 0.015 |
| BON | 0.271 | 0.242 | 0.115 | -0.010 | 0.032 | 0.508 | 0.026 |
| CAD | 0.233 | 0.309 | 0.110 | 0.087 | 0.137 | 0.443 | 0.030 |
| CAG | 0.217 | 0.345 | 0.379 | -0.020 | 0.176 | 0.430 | 0.016 |
| CAR | 0.264 | 0.409 | 0.161 | -0.050 | 0.078 | 0.544 | 0.032 |
| CAS | 0.227 | 0.303 | 0.101 | 0.029 | 0.162 | 0.478 | 0.030 |
| CAZ | 0.215 | 0.216 | 0.133 | -0.230 | 0.120 | 0.468 | 0.030 |
| CEN | 0.266 | 0.319 | 0.126 | -0.010 | 0.028 | 0.480 | 0.049 |
| COC | 0.246 | 0.372 | 0.158 | -0.010 | 0.023 | 0.484 | 0.033 |
| COD | 0.227 | 0.229 | 0.102 | -0.010 | 0.077 | 0.442 | 0.022 |
| CPA | 0.249 | 0.315 | 0.167 | -0.020 | 0.039 | 0.472 | 0.019 |
| CUE | 0.240 | 0.341 | 0.144 | -0.010 | 0.029 | 0.462 | 0.034 |
| EST | 0.233 | 0.206 | 0.143 | -0.010 | 0.167 | 0.492 | 0.052 |
| FUE | 0.211 | 0.264 | 0.132 | -0.020 | 0.177 | 0.471 | 0.050 |
| GAU | 0.231 | 0.194 | 0.135 | 0.035 | 0.183 | 0.499 | 0.032 |
| GEA | 0.239 | 0.211 | 0.119 | -0.070 | 0.069 | 0.476 | 0.023 |
| GIR | 0.171 | 0.285 | 0.294 | -0.070 | 0.294 | 0.432 | 0.020 |

| | | | | | | | |
|---|---|---|---|---|---|---|---|
| GUA | 0.204 | 0.312 | 0.348 | -0.070 | 0.227 | 0.437 | 0.014 |
| HOU | 0.227 | 0.310 | 0.141 | -0.010 | 0.107 | 0.463 | 0.032 |
| JUB | 0.216 | 0.173 | 0.119 | -0.120 | 0.217 | 0.487 | 0.035 |
| KUD | 0.185 | 0.134 | 0.108 | -0.220 | 0.353 | 0.422 | 0.042 |
| LAC | 0.235 | 0.298 | 0.137 | -0.020 | 0.109 | 0.473 | 0.053 |
| LAM | 0.226 | 0.304 | 0.099 | -0.010 | 0.157 | 0.452 | 0.027 |
| LCO | 0.226 | 0.248 | 0.122 | -0.020 | 0.116 | 0.478 | 0.027 |
| LEI | 0.227 | 0.256 | 0.082 | -0.020 | 0.077 | 0.460 | 0.029 |
| LIM | 0.232 | 0.292 | 0.136 | -0.020 | 0.114 | 0.474 | 0.058 |
| MAD | 0.202 | 0.157 | 0.132 | -0.040 | 0.310 | 0.495 | 0.026 |
| MAG | 0.189 | 0.259 | 0.322 | 0.007 | 0.306 | 0.449 | 0.016 |
| MAU | 0.221 | 0.210 | 0.336 | -0.020 | 0.219 | 0.446 | 0.019 |
| MAZ | 0.236 | 0.191 | 0.077 | -0.010 | 0.039 | 0.460 | 0.032 |
| MIM | 0.233 | 0.322 | 0.153 | 0.024 | 0.102 | 0.457 | 0.061 |
| MON | 0.219 | 0.278 | 0.376 | 0.033 | 0.286 | 0.469 | 0.014 |
| MTG | 0.258 | 0.267 | 0.186 | 0.011 | 0.063 | 0.452 | 0.015 |
| MUR | 0.203 | 0.272 | 0.349 | -0.050 | 0.288 | 0.474 | 0.023 |
| OLB | 0.245 | 0.268 | 0.144 | 0.008 | 0.034 | 0.453 | 0.017 |
| OLO | 0.224 | 0.296 | 0.150 | 0.013 | 0.112 | 0.459 | 0.070 |
| ONA | 0.231 | 0.275 | 0.117 | -0.030 | 0.075 | 0.474 | 0.017 |
| ORI | 0.245 | 0.267 | 0.190 | -0.010 | 0.089 | 0.483 | 0.020 |
| PAN | 0.164 | 0.281 | 0.242 | -0.040 | 0.376 | 0.445 | 0.049 |
| PCI | 0.209 | 0.291 | 0.145 | 0.135 | 0.202 | 0.493 | 0.059 |
| PDL | 0.233 | 0.267 | 0.150 | 0.036 | 0.089 | 0.469 | 0.022 |
| PET | 0.234 | 0.334 | 0.161 | -0.010 | 0.098 | 0.465 | 0.058 |
| PEZ | 0.254 | 0.266 | 0.184 | -0.040 | 0.081 | 0.482 | 0.028 |
| PFQ | 0.224 | 0.328 | 0.132 | -0.020 | 0.100 | 0.447 | 0.031 |
| PIN | 0.204 | 0.354 | 0.383 | 0.041 | 0.235 | 0.451 | 0.015 |
| PLE | 0.231 | 0.338 | 0.153 | -0.040 | 0.098 | 0.467 | 0.040 |
| PUE | 0.231 | 0.390 | 0.147 | -0.060 | 0.145 | 0.463 | 0.024 |
| QUA | 0.242 | 0.275 | 0.147 | -0.010 | 0.058 | 0.466 | 0.023 |
| QUI | 0.226 | 0.266 | 0.104 | -0.090 | 0.051 | 0.441 | 0.024 |
| RIO | 0.256 | 0.277 | 0.175 | -0.020 | 0.053 | 0.507 | 0.045 |
| ROS | 0.223 | 0.294 | 0.383 | -0.020 | 0.183 | 0.464 | 0.025 |
| SAC | 0.228 | 0.365 | 0.134 | -0.030 | 0.105 | 0.421 | 0.048 |
| SAH | 0.238 | 0.329 | 0.130 | -0.030 | 0.032 | 0.475 | 0.028 |
| SAL | 0.245 | 0.292 | 0.112 | -0.010 | 0.038 | 0.454 | 0.025 |
| SCD | 0.225 | 0.246 | 0.135 | -0.060 | 0.080 | 0.463 | 0.019 |
| SEB | 0.208 | 0.250 | 0.318 | -0.030 | 0.176 | 0.428 | 0.021 |
| SEG | 0.216 | 0.317 | 0.116 | -0.060 | 0.110 | 0.469 | 0.079 |
| SID | 0.114 | 0.094 | 0.065 | -0.160 | 0.640 | 0.487 | 0.018 |
| SIE | 0.225 | 0.373 | 0.132 | 0.004 | 0.130 | 0.422 | 0.031 |
| SIN | 0.230 | 0.248 | 0.118 | -0.070 | 0.086 | 0.490 | 0.023 |
| SOB | 0.236 | 0.278 | 0.116 | 0.050 | 0.060 | 0.496 | 0.029 |
| STJ | 0.225 | 0.318 | 0.153 | 0.000 | 0.112 | 0.466 | 0.053 |
| TAJ | 0.177 | 0.132 | 0.098 | 0.034 | 0.442 | 0.461 | 0.019 |
| TAL | 0.230 | 0.320 | 0.133 | -0.040 | 0.068 | 0.474 | 0.056 |
| TAM | 0.185 | 0.132 | 0.134 | -0.130 | 0.389 | 0.451 | 0.027 |

| | | | | | | | |
|---|---|---|---|---|---|---|---|
| TBY | 0.225 | 0.257 | 0.089 | 0.012 | 0.082 | 0.485 | 0.013 |
| TOC | 0.197 | 0.289 | 0.336 | -0.050 | 0.287 | 0.458 | 0.020 |
| VER | 0.223 | 0.313 | 0.146 | 0.034 | 0.110 | 0.480 | 0.022 |
| VMA | 0.229 | 0.230 | 0.115 | -0.060 | 0.087 | 0.471 | 0.017 |
| VMQ | 0.257 | 0.341 | 0.161 | -0.070 | 0.029 | 0.447 | 0.054 |

**Table S4.** Climatic variables used in the genomics offset analyses, as provided by the Climate Downscaling Tool (ClimateDT, https://www.ibbr.cnr.it/climate-dt/).

| Label | Description | Unit |
|---|---|---|
| bio1 | Mean annual temperature | Celsius degrees (°C) |
| bio3 | Isothermality (bio2/bio7) (×100) | Index |
| bio4 | Temperature seasonality (standard deviation ×100) | Celsius degrees (°C) |
| bio12 | Annual precipitation | Millimeters (mm) |
| bio15 | Precipitation seasonality (coefficient of variation) | Index |
| SHM | Summer Heat Moisture index | °C/mm |

**Table S5**. Models a) without and b) with gene pool effect as a random factor, evaluating the effect of population marginality on population-specific Jost's *D* (average pairwise values), a measure of genetic differentiation that is independent of levels of genetic diversity. Significant fixed effects are given with their associated point estimates and 95% confidence intervals [in brackets]. $R^2$: variance explained by fixed factors or both fixed (marginal $R^2$) and random (conditional $R^2$) factors; ns: not significant.

a) Models without gene pool effect.

| Model [$R^2$] | Genetic indicator | Fixed-effect #1 | Estimate [95% CIs] | Fixed-effect #2 | Estimate [95% CIs] | Fixed-effect #3 | Estimate [95% CIs] | Interaction | Estimate [95% CIs] |
|---|---|---|---|---|---|---|---|---|---|
| M5bis [0.61] | *Population-specific Jost's D* | Centroid | 0.76 [0.44, 1.09] | Second nearest-core | 1.88 [0.94, 2.82] | North-South | -2.37 [-3.29, -1.44] | Second nearest-core × North-South | -2.03 [-2.95, -1.12] |

b) Models with gene pool effect as random factor.

| Model [$R^2$] | Genetic indicator | Fixed-effect #1 | Estimate [95% CIs] | Fixed-effect #2 | Estimate [95% CIs] | Interaction | Estimate [95% CIs] |
|---|---|---|---|---|---|---|---|
| M12bis [0.31, 0.90] | *Population-specific Jost's D* | Centroid | 0.88 [0.32, 1.44] | North-South | -1.76 [-2.57, -0.94] | Centroid × North-South | -1.19 [-1.73, -0.66] |

**Figure S1**. Distribution range of *P. pinaster* considering only natural populations (i.e., excluding plantations). This map was produced based on the distribution map of *P. pinaster* by Caudullo et al. (2017), which contains both natural populations and plantations. The boundaries of *P. pinaster* natural distribution were delimited with QGIS v3.22 based on information available in Alía et al., 1996; Marques et al., 2012; Abad-Viñas et al., 2016; Fkiri et al., 2019 and Wahid et al., 2004, 2006 and taking into account the National Forest Inventories of Spain, Italy and France.

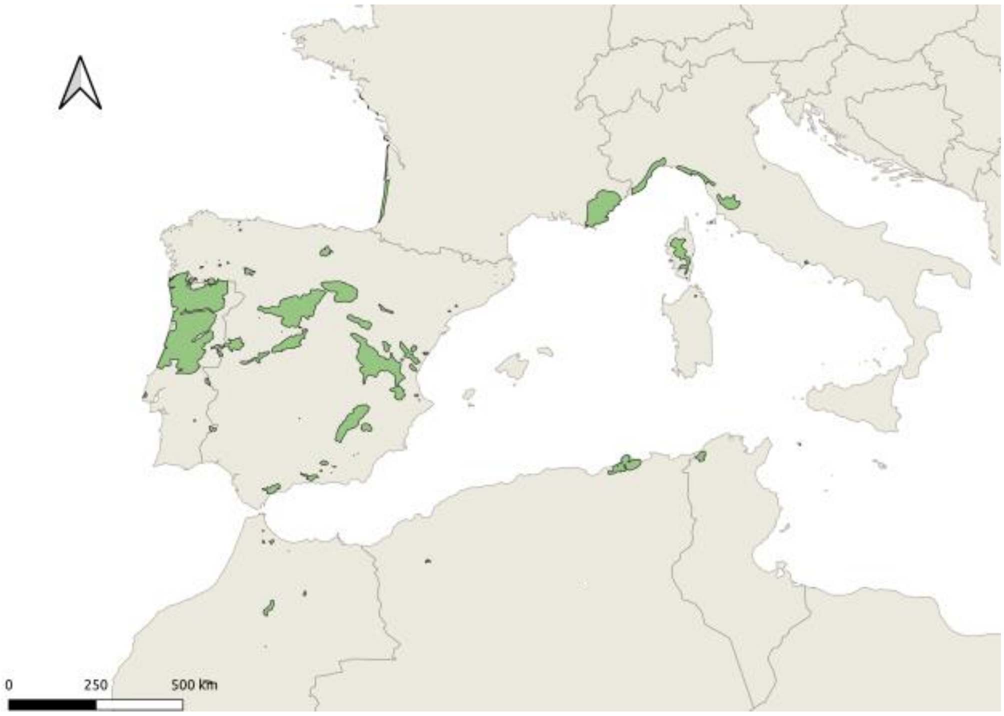

**Figure S2.** Map showing the position of the centroid of the maritime pine natural distribution (in blue), which corresponds to the location whose geographic coordinates are the average of all locations where the species is naturally present (see Figure S1).

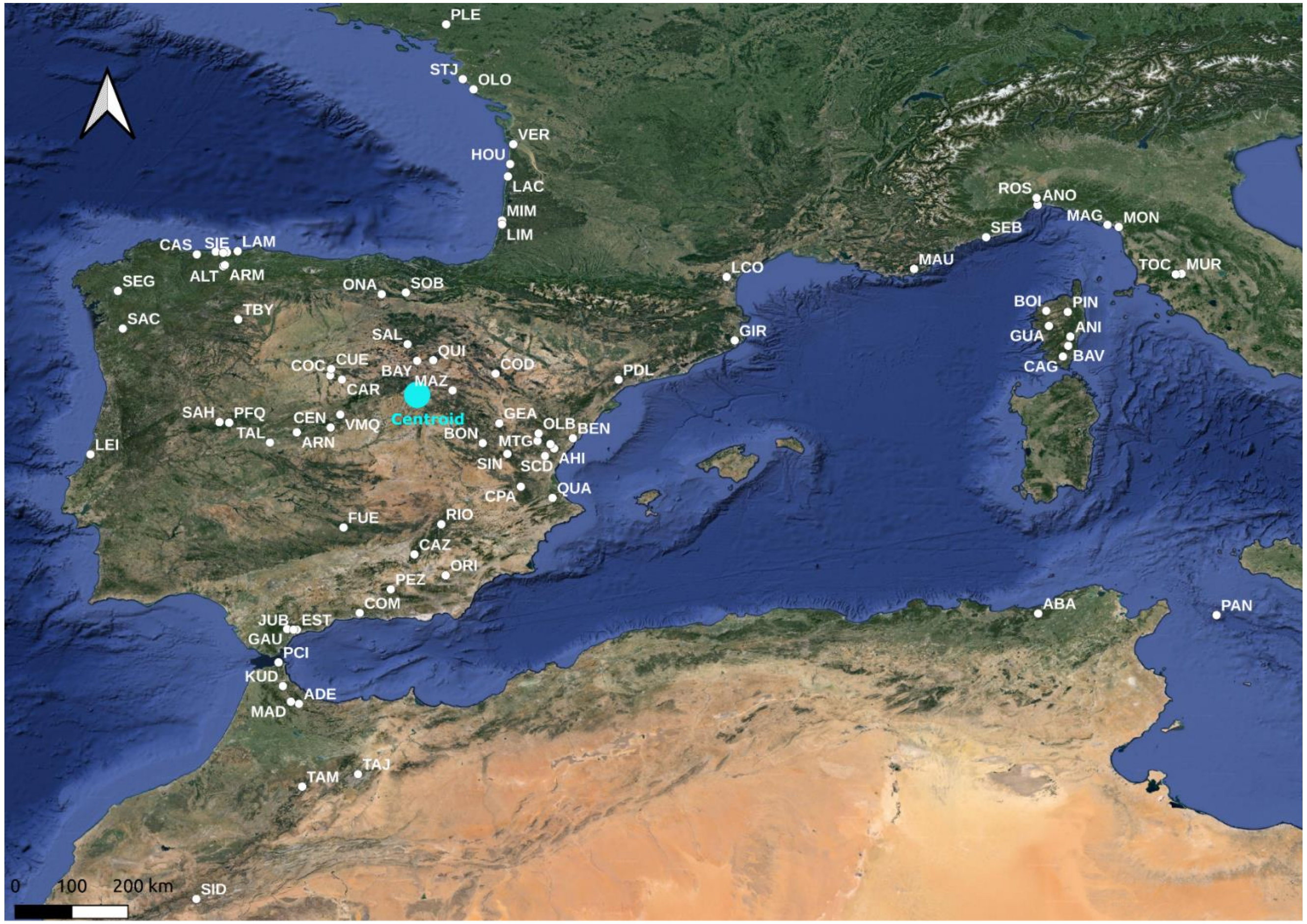

**Figure S3**. Correlation of standardized marginality indices; a) Pairwise correlogram based on Pearson's correlation coefficients; b) Plot of the two first axes (explaining 54.3% of the variance) of the Principal Component Analysis (PCA).

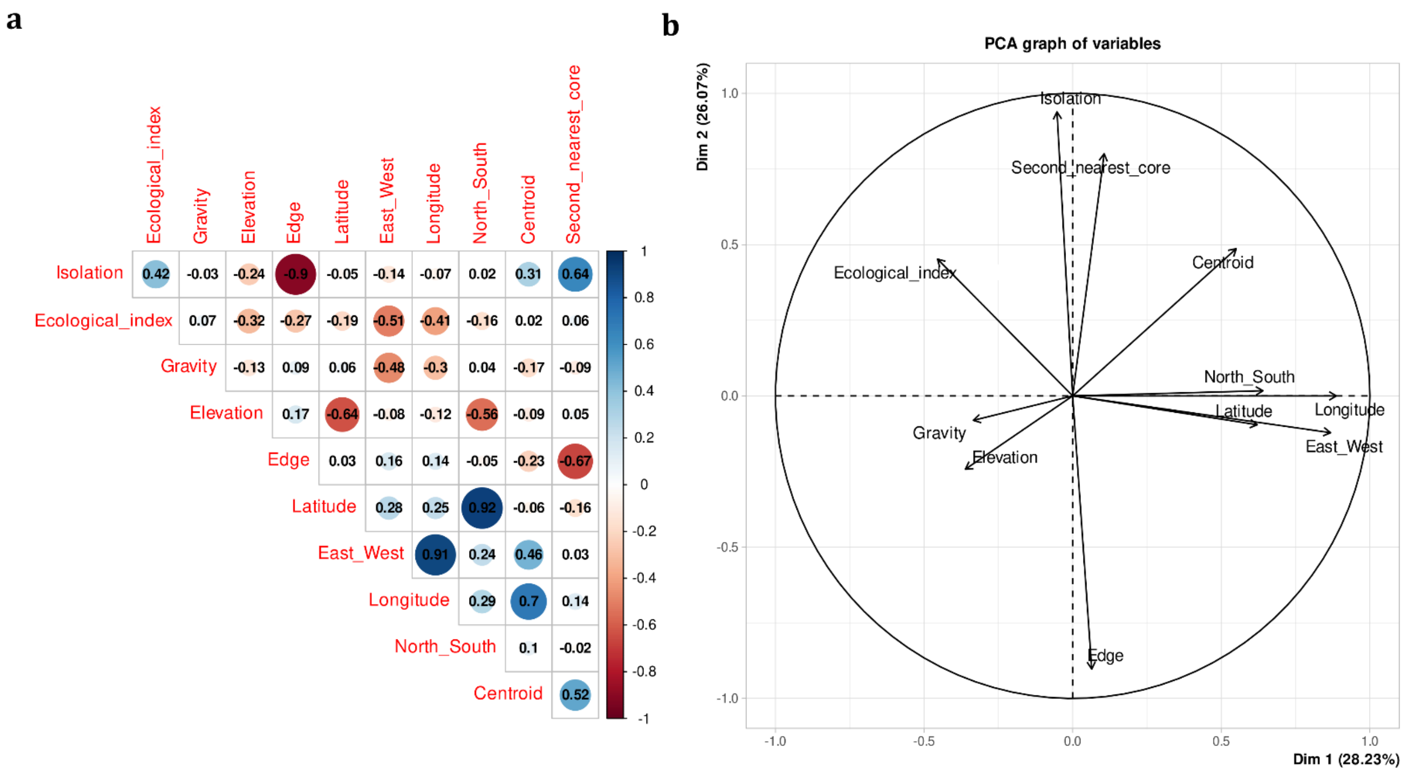

**Figure S4**. Correlation of genetic indicator estimates using the full dataset (10,185 SNPs) with missing data lower than 30% and a dataset (6,390 SNPs) with missing data lower than 5%, for a) overall genetic diversity, $F_{IS}$, population-specific $F_{ST}$, and genomic offset; and b) linear regression between recessive genetic load (RGL) calculated at the individual level from the full dataset with missing data lower than 30% (RGL_miss30) and a dataset with missing data lower than 5% (RGL_miss5). Numbers in a) correspond to pairwise Pearson's correlation coefficients. See full description of statistics in Table S3 and/or in the main text.

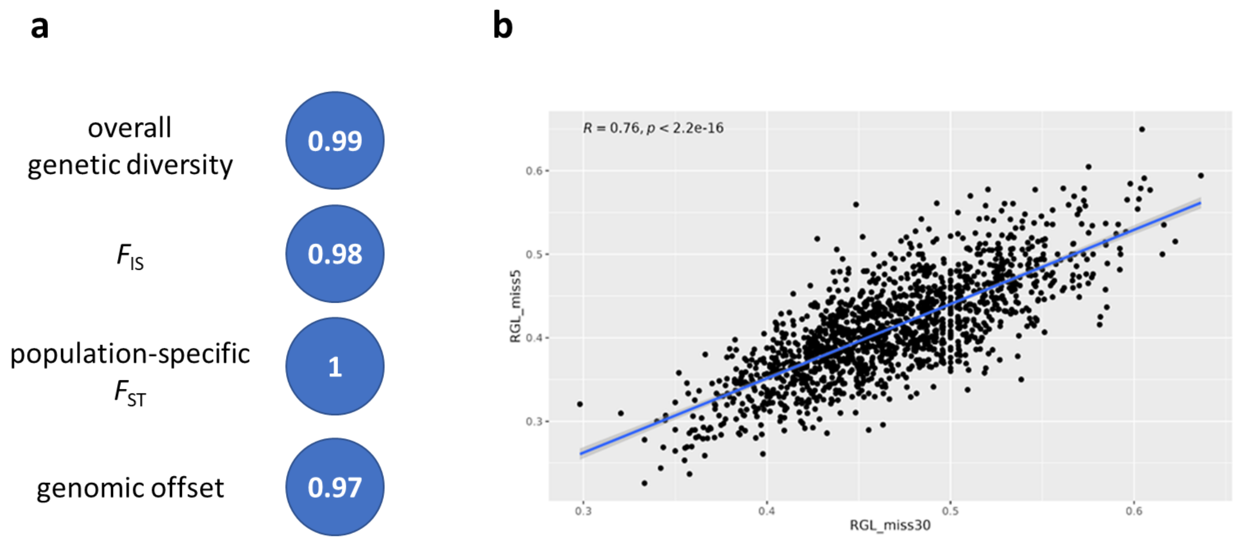

**Figure S5.** Pairwise correlogram based on Pearson's correlation coefficients of genomic offset predictions for the five GCMs used in this study.

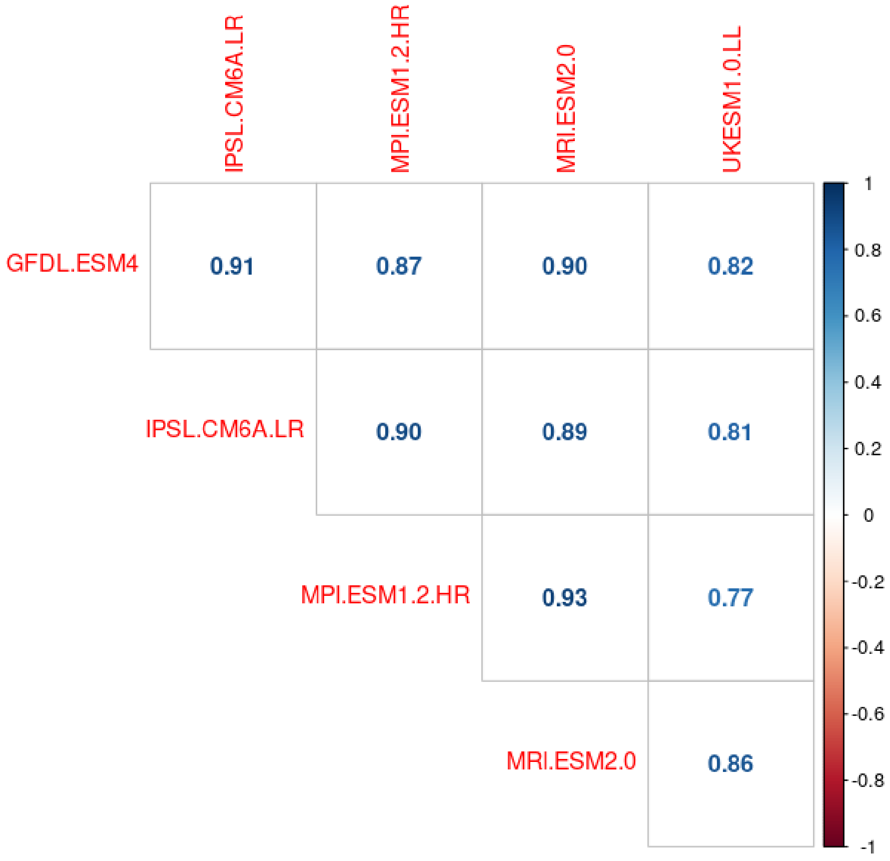

**Figure S6.** Maps depicting the geographic distribution of the marginality indices selected in our study for *P. pinaster* populations. a) Centroid; b) Second nearest-core; c) Gravity; d) North-South; and e) Ecological index.

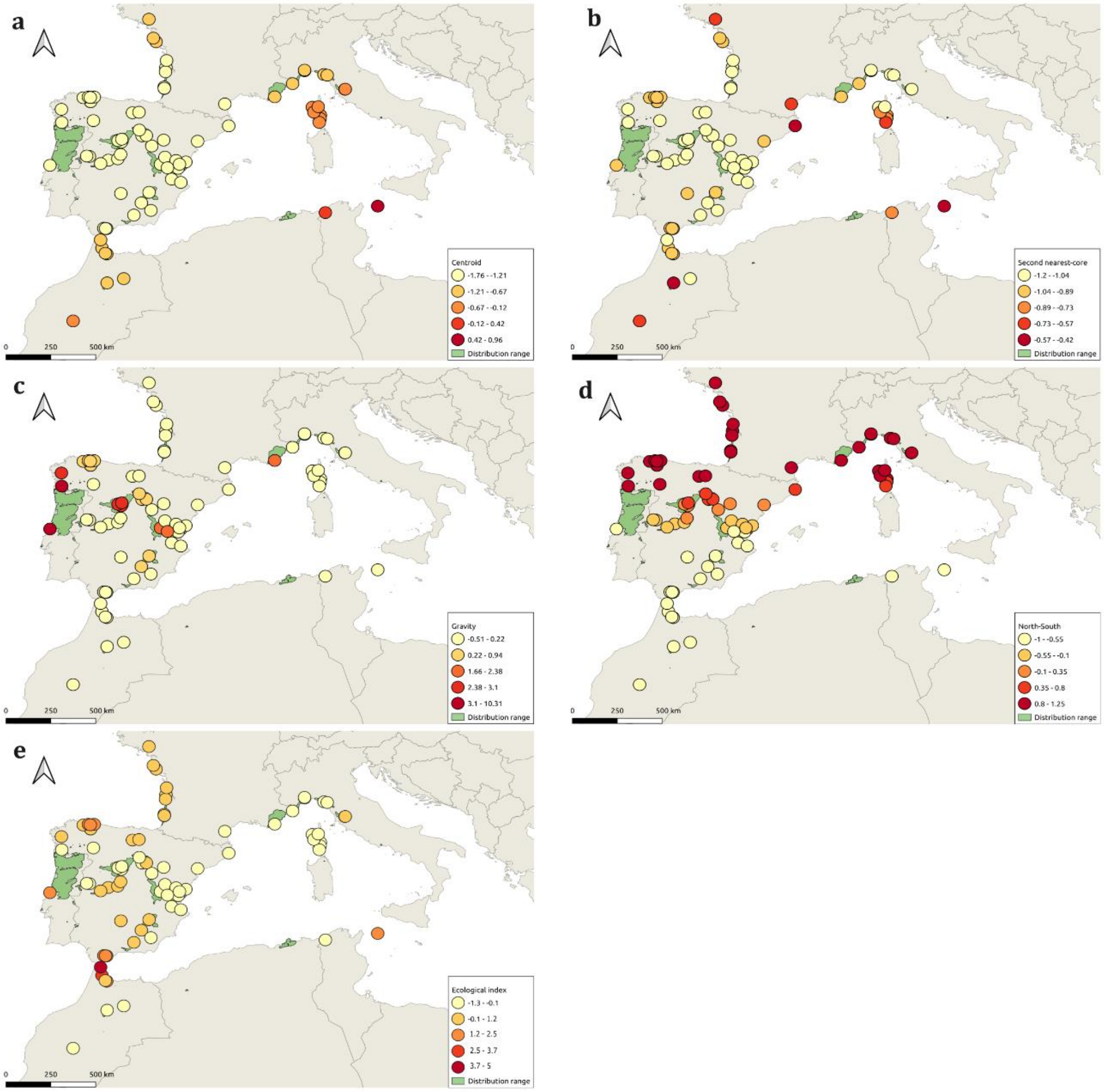

**Figure S7**. Breakdown of interaction effects between the Second nearest-core and the North-South indices in M1 (Overall genetic diversity; 1-$Q_{inter}$ corrected for sample size) and M5 (Genetic differentiation; population-specific $F_{ST}$). The fitted interaction values were calculated using the *effects* R package and the visual representation was produced with *ggplot2* R package. These figures show the decomposition of the interaction between distance to the second-nearest core (distance increases from left to right) and the latitudinal position of the populations (North-South index; where -1 represent southern latitudes and 1 represents northern latitudes), for a) overall genetic diversity and b) genetic differentiation.

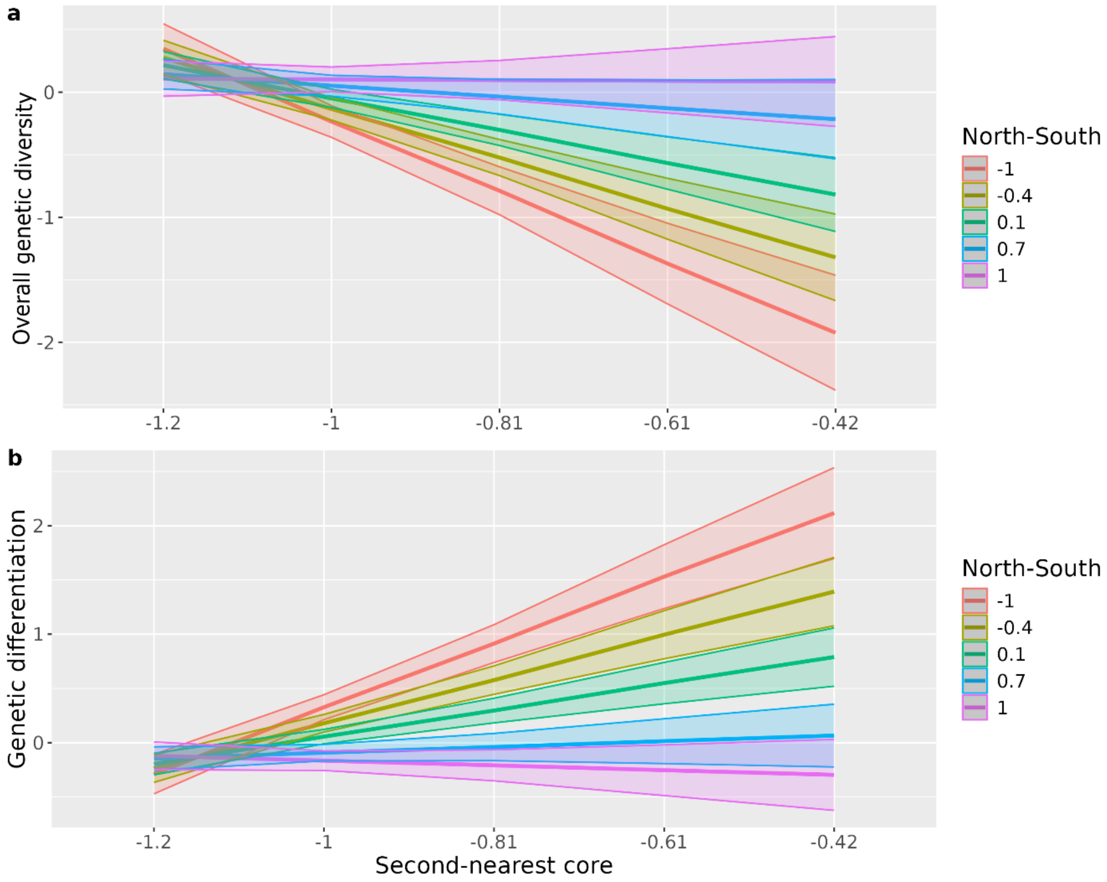

**Figure S8.** Map representing the geographical distribution of a) Genetic diversity for GEA outliers, b) Genetic diversity for general outliers, c) Population inbreeding ($F_{IS}$) and d) Recessive genetic load, for *P. pinaster* populations across its distribution range.

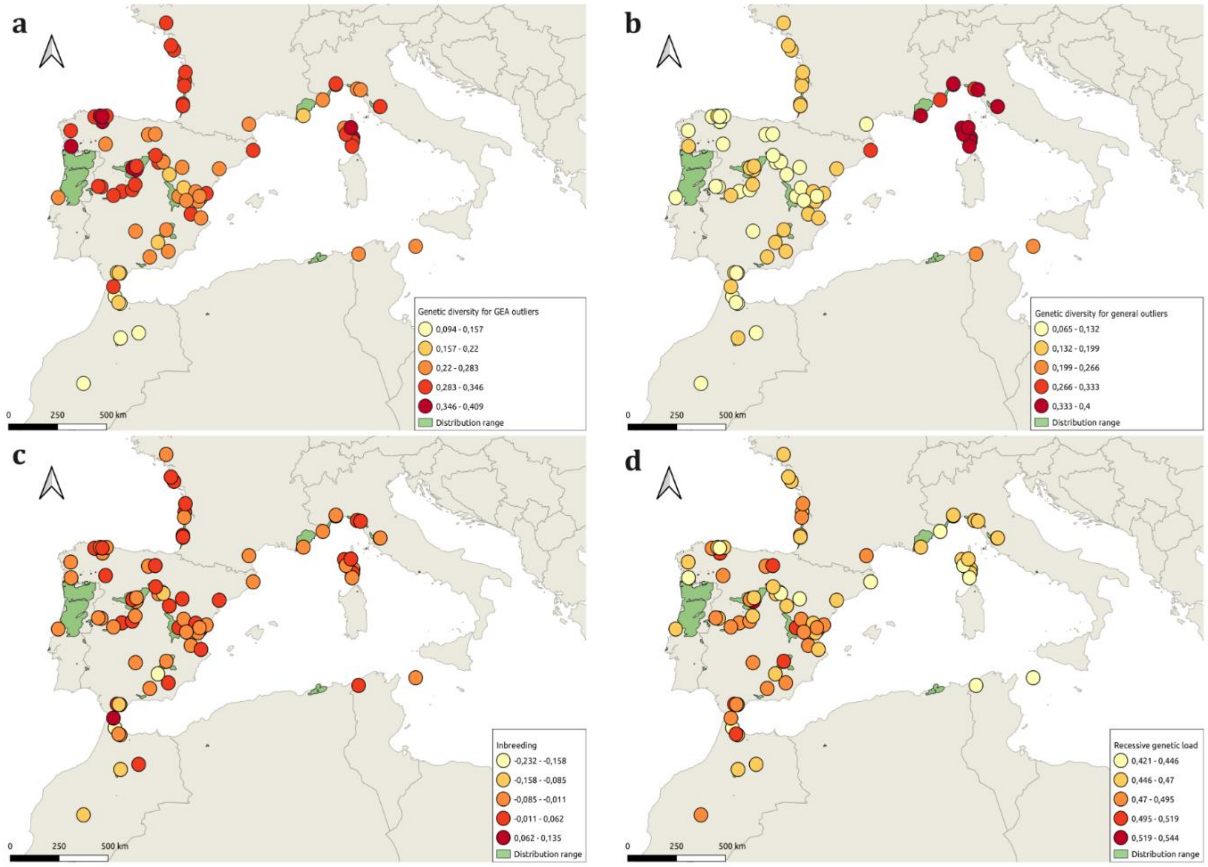

**Figure S9**. Breakdown of interaction effects between the Centroid and the North-South indices in M3 (Genetic diversity based on general outliers). The fitted interaction values were calculated using the *effects* R package and the visual representation was produced with *ggplot2* R package. This figure shows the decomposition of the interaction between distance to the centroid (distance increases from lef t to right) and the latitudinal position of the populations (North-South index; where -1 represent southern latitudes and 1 represents northern latitudes).

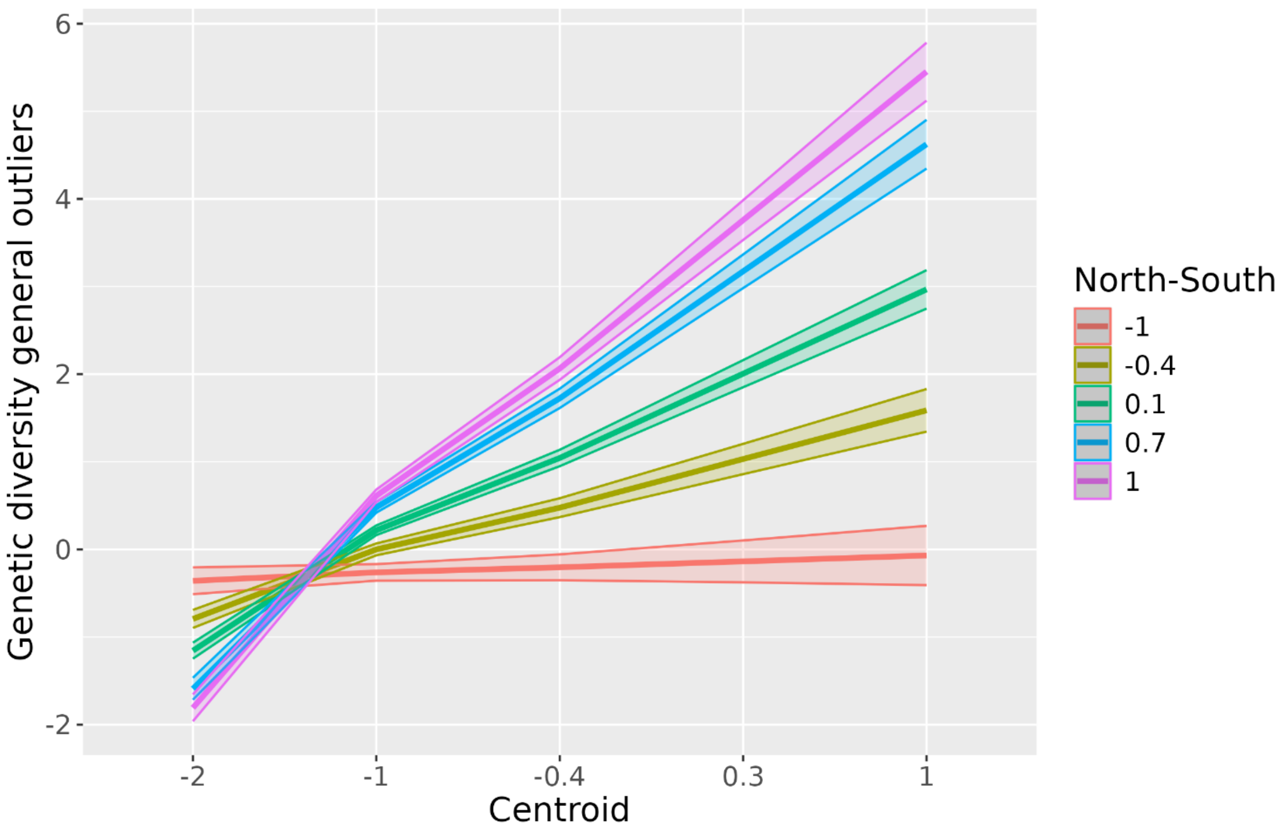

**Figure S10**. Breakdown of interaction effects between the Centroid and the North-South indices in M8 (Overall genetic diversity; 1-$Q_{inter}$ corrected for sample size) and M12 (Genetic differentiation; population-specific $F_{ST}$). The fitted interaction values were calculated using the *effects* R package and the visual representation was produced with *ggplot2* R package. These figures show the decomposition of the interaction between distance to the second-nearest core (distance increases from left to right) and the latitudinal position of the populations (North-South index; where -1 represent southern latitudes and 1 represents northern latitudes), for a) overall genetic diversity and b) genetic differentiation.

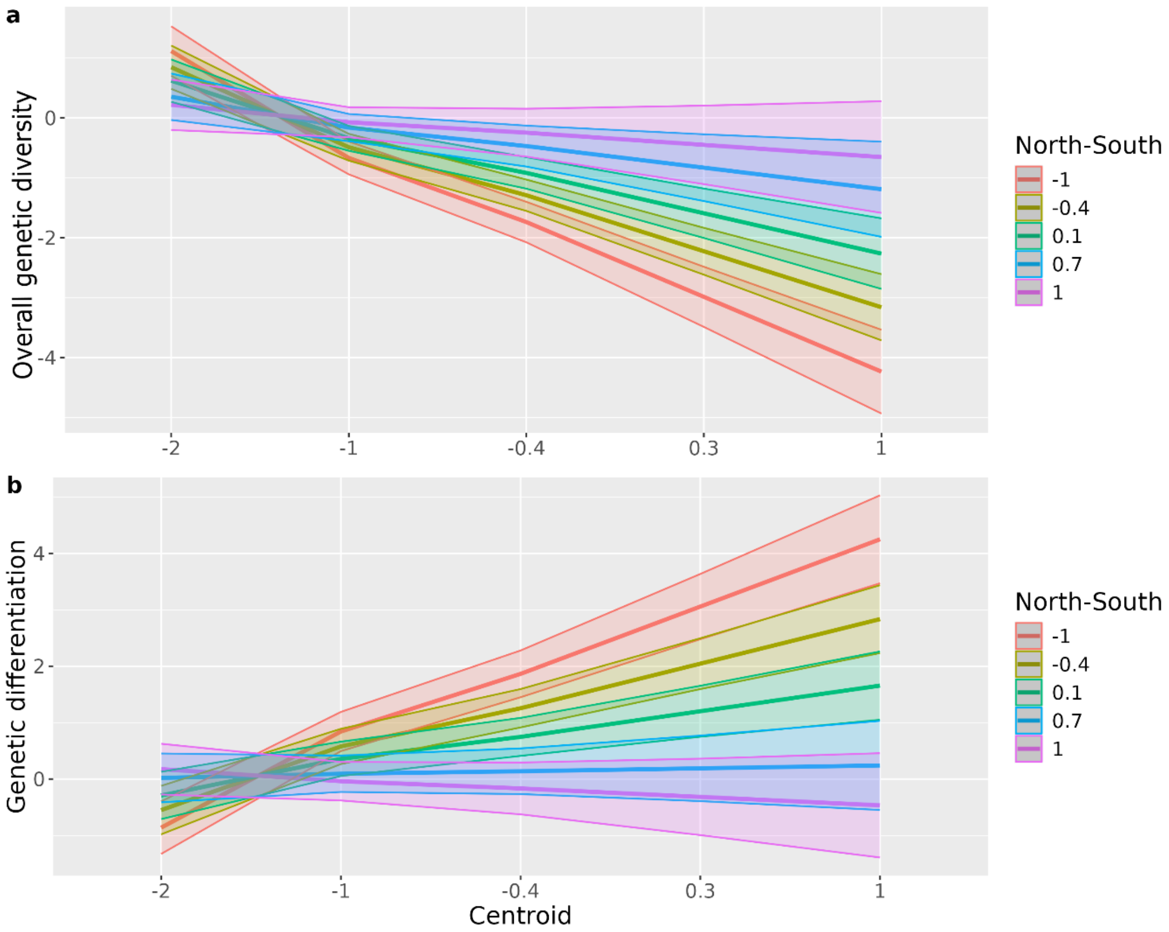

**Figure S11.** Gradient Forest (GF) analyses: a) Turnover functions and b) Mean accuracy importance, for each climate covariate (similar results were obtained for the mean importance weighted by SNP $R^2$). The full names and units of the climate covariates are given in Table S4. Find also additional details at https://anonymous.4open.science/r/ReadyToGO_Pinpin-FA56/README.md.

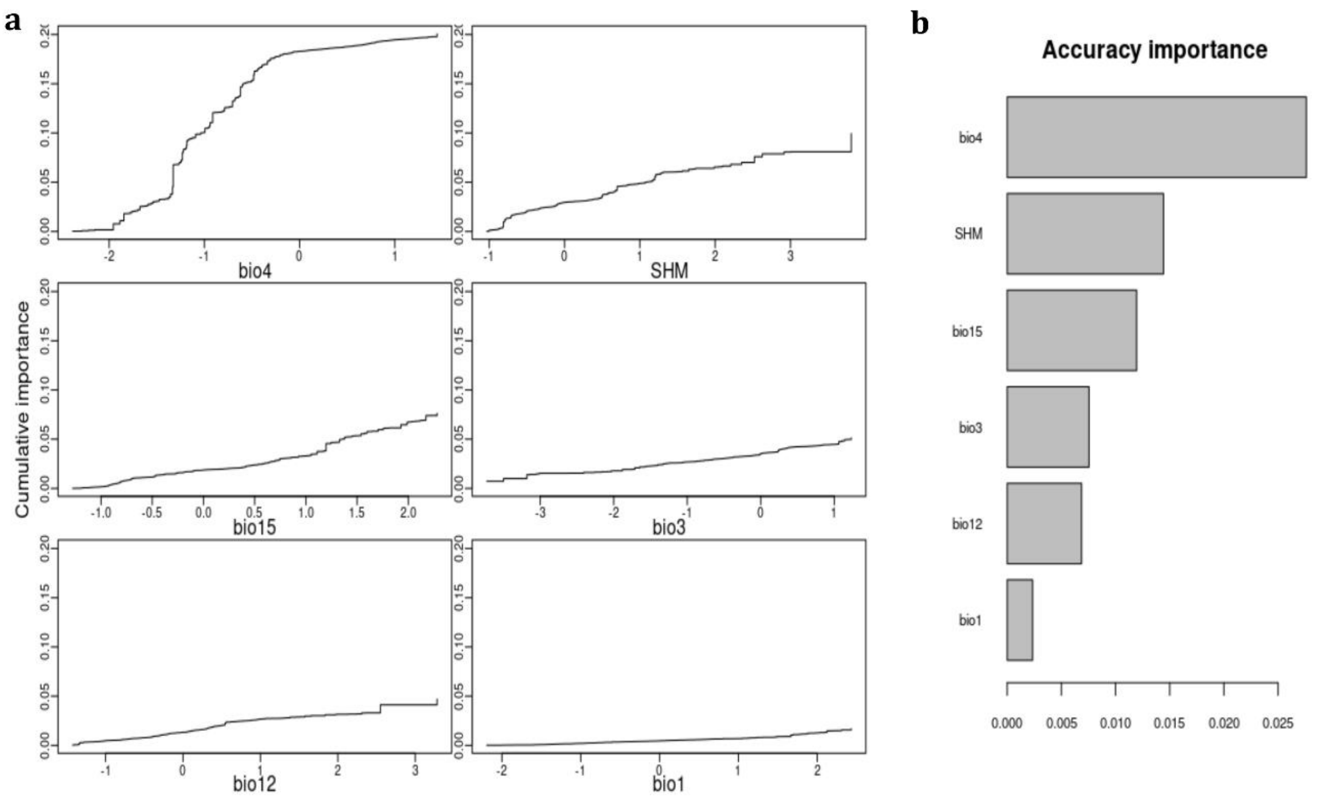

**Figure S12.** Diagnostic plots to check model fit to data. QQ-plots a, c, e) and residual plots b, d, f) were computed for M1, M2 and M3, respectively.

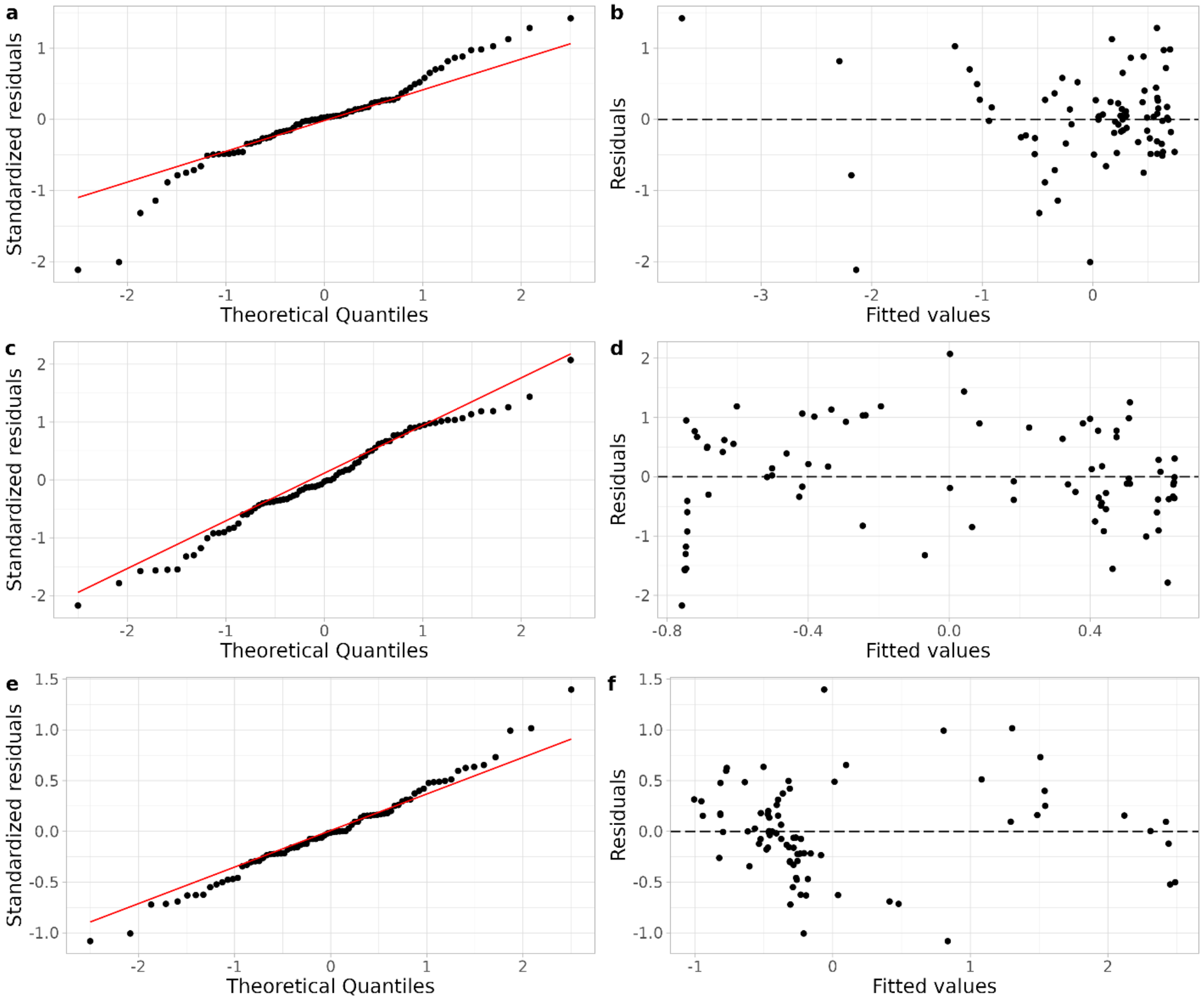

**Figure S13.** Diagnostic plots to check model fit to data. QQ-plots a, c) and residual plots b, d) were computed for M8 and M9, respectively.

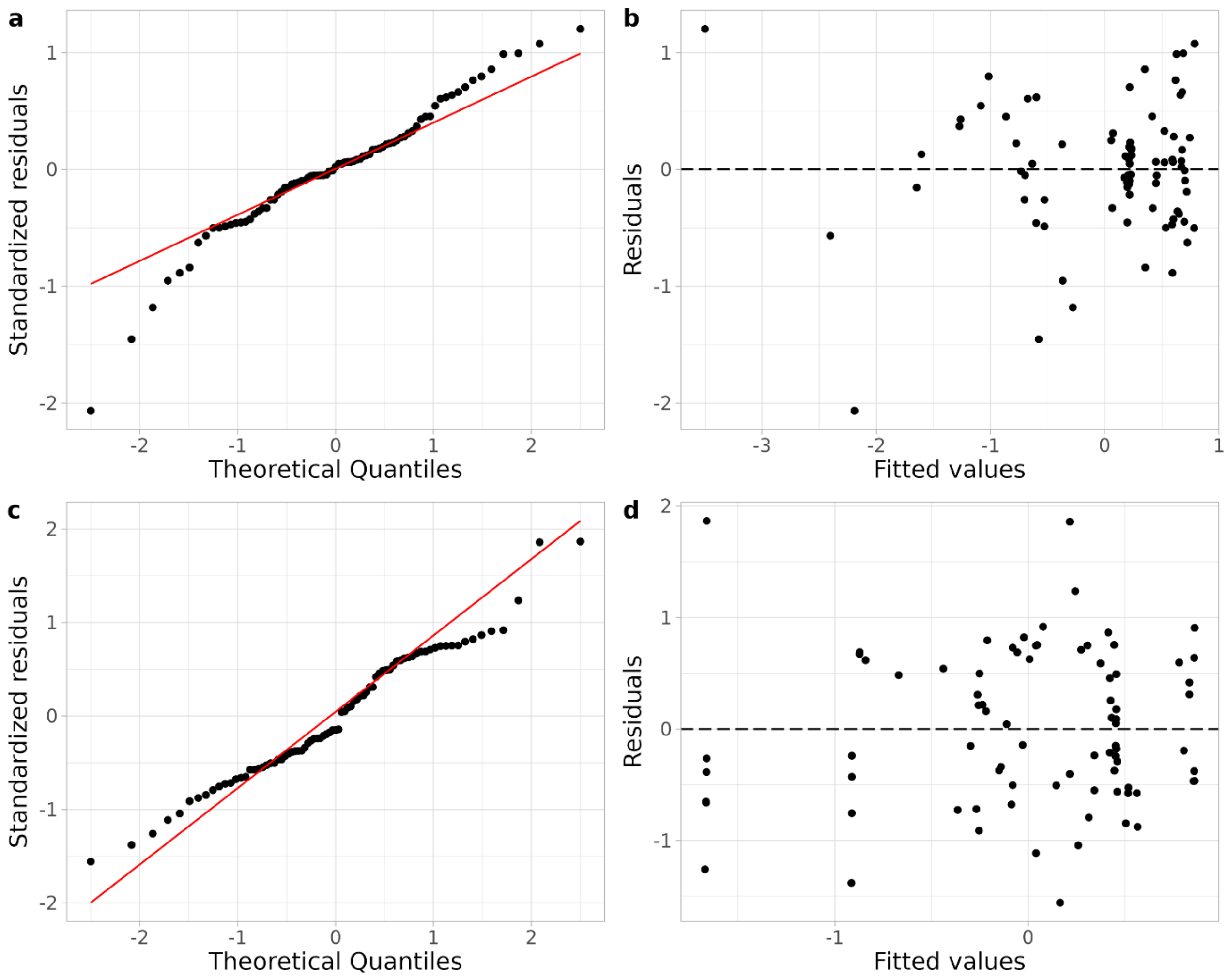

**Figure S14.** Diagnostic plots to check model fit to data. QQ-plots a, c) and residual plots b, d) were computed for M5 and M5bis, respectively.

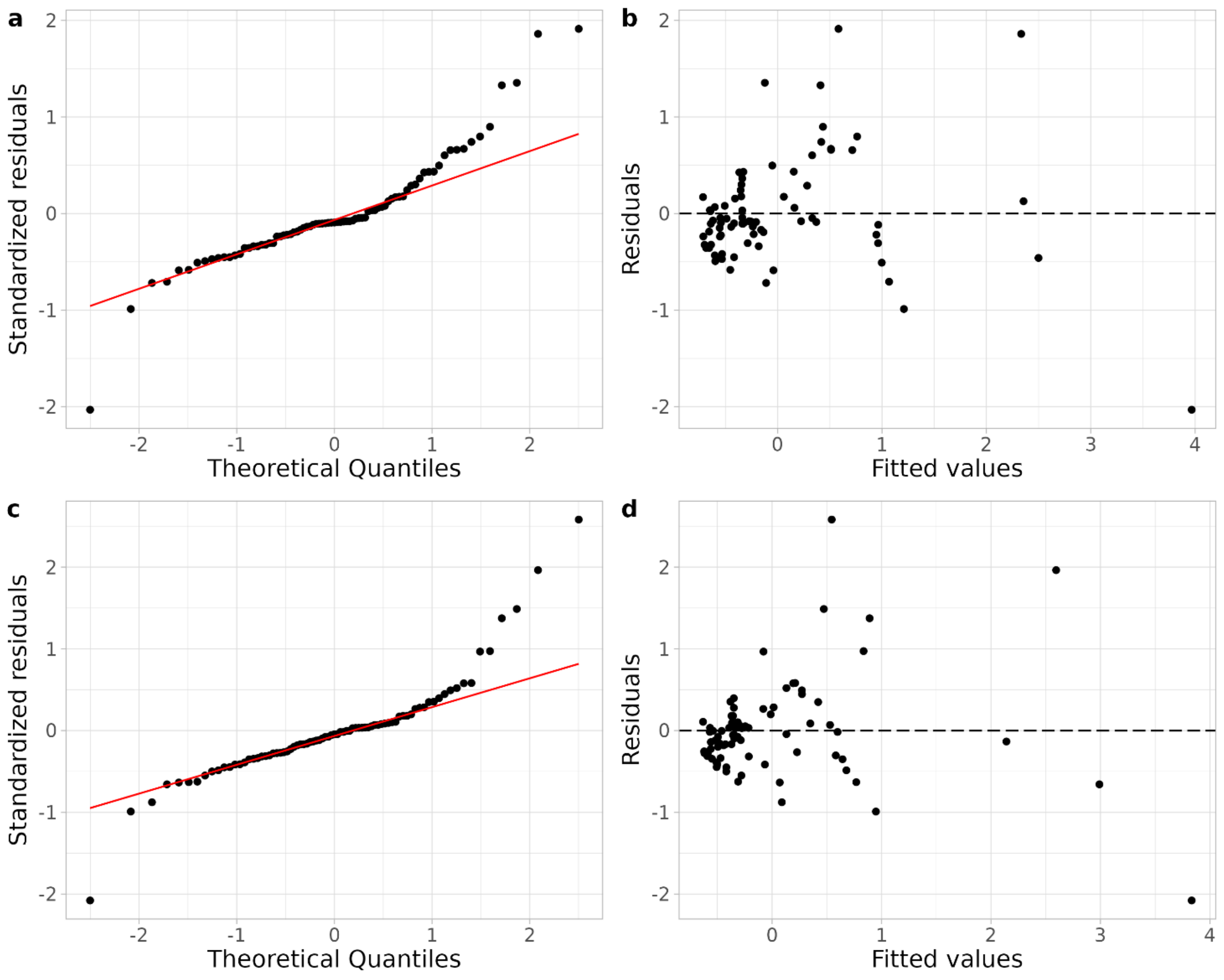

**Figure S15.** Diagnostic plots to check model fit to data. QQ-plots a, c) and residual plots b, d) were computed for M12 and M12bis, respectively.

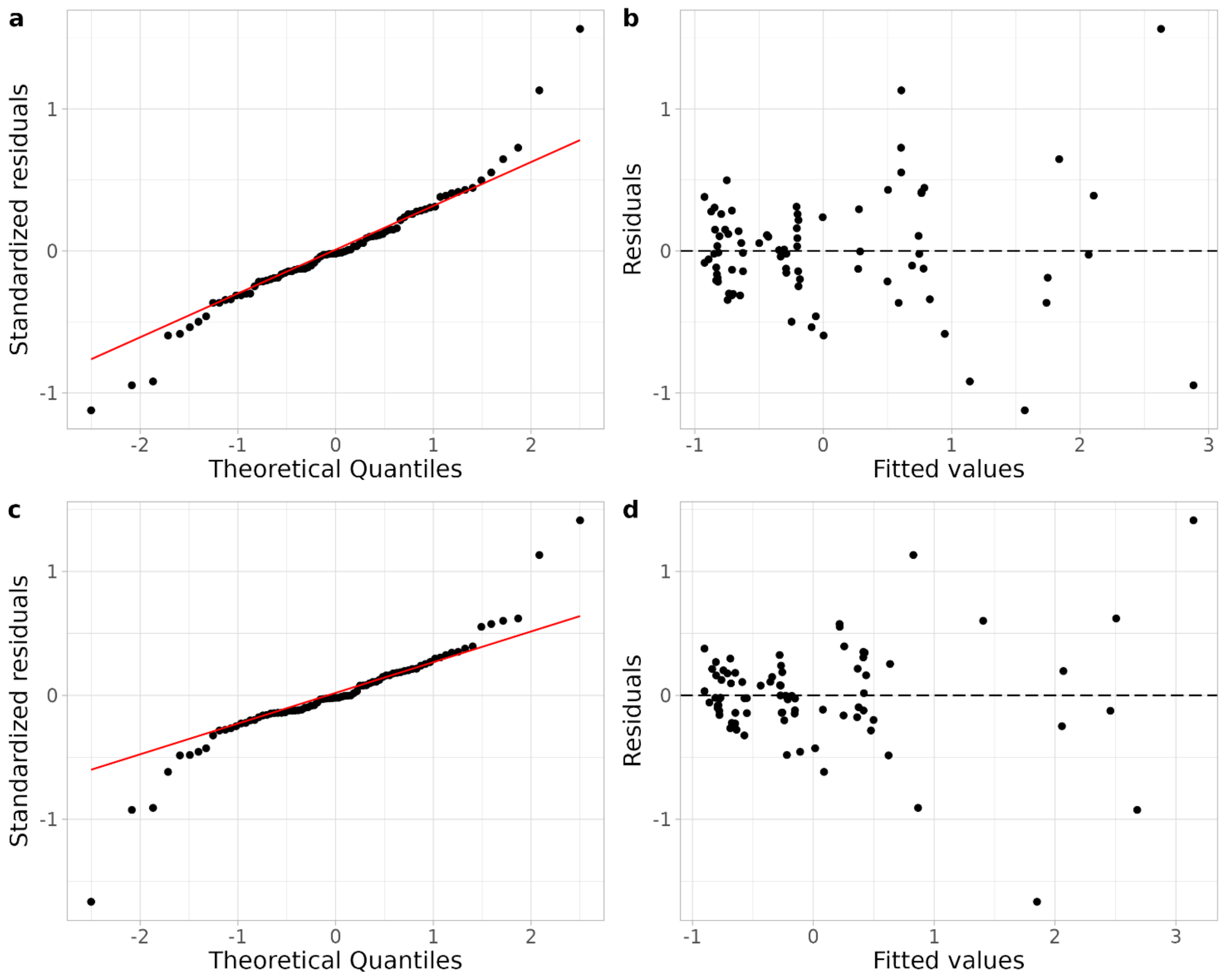

**Figure S16.** Diagnostic plots to check model fit to data. QQ-plots a, c) and residual plots b, d) were computed for M6 and M7, respectively.

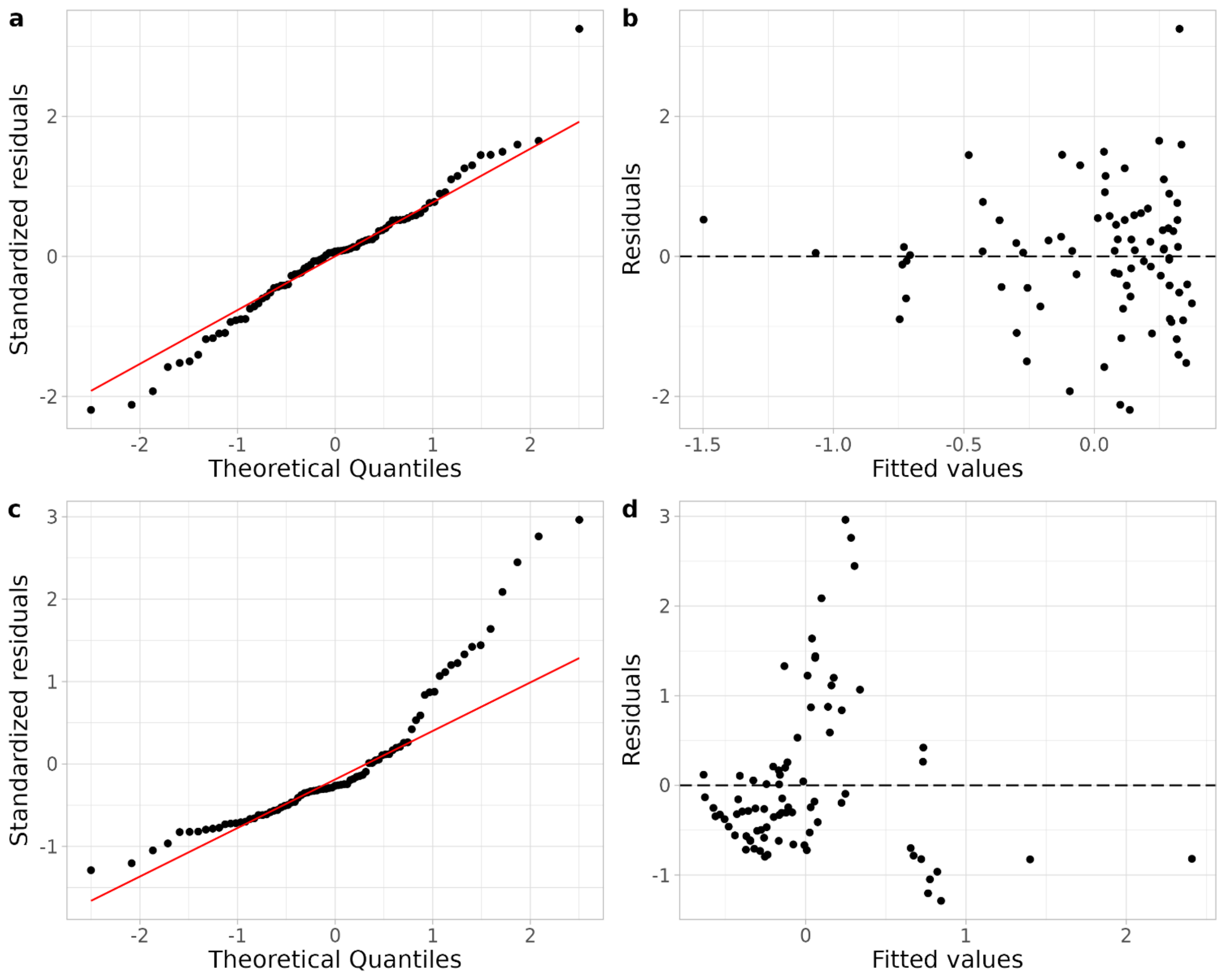

**Figure S17.** Diagnostic plots to check model fit to data. QQ-plots a) and residual plots b) were computed for M14, respectively.

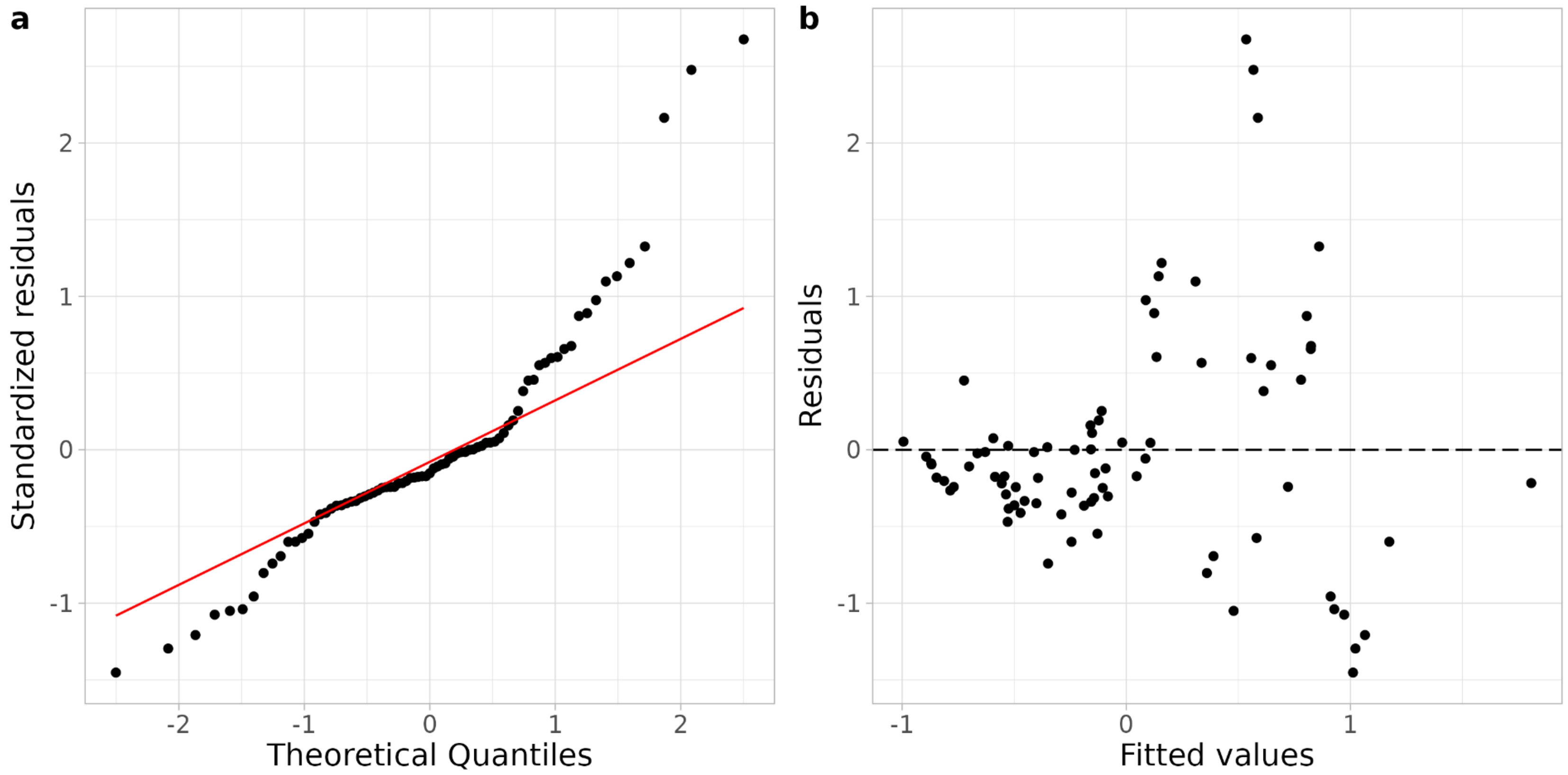
